# Life history genomic regions explain differences in Atlantic salmon marine diet specialization

**DOI:** 10.1101/754440

**Authors:** Tutku Aykanat, Martin Rasmussen, Mikhail Ozerov, Eero Niemelä, Lars Paulin, Juha-Pekka Vähä, Kjetil Hindar, Vidar Wennevik, Torstein Pedersen, Martin-A. Svenning, Craig R. Primmer

## Abstract

1. Animals employ various foraging strategies along their ontogeny to acquire energy, and with varying degree of efficiencies, to support growth, maturation and subsequent reproduction events. Individuals that can efficiently acquire energy early are more likely to mature at an earlier age, as a result of faster energy gain which can fuel maturation and reproduction.
2. We aimed to test the hypothesis that heritable resource acquisition variation that co-varies with efficiency along the ontogeny would influence maturation timing of individuals.
3. To test this hypothesis, we utilized Atlantic salmon as a model which exhibit a simple, hence trackable, genetic control of maturation age. We then monitored the variation in diet acquisition (quantified as the stomach fullness and composition) of individuals with different ages, and linked it genomic regions (haploblocks) that were previously identified to be associated with age-at-maturity.
4. Consistent with the hypothesis, we demonstrated that one of the life history genomic regions tested (*six6*) was indeed associated with age-dependent differences in stomach fullness. Prey composition was marginally linked to both genomic regions (*six6* and *vgll3*). We further showed Atlantic salmon switched to the so-called “feast and famine” strategy along the ontogeny, where older age groups exhibited heavier stomach content, but that came at the expense of running on empty more often.
5. These results suggest genetic variation underlying resource utilization variation may explain the genetic basis of age structure in Atlantic salmon. Given that ontogenetic diet has a genetic component and the strong spatial diversity associated with these genomic regions, we predict populations with diverse maturation age will have diverse evolutionary responses to future changes in marine food-web structures.

## Introduction

Diet acquisition is a strong evolutionary force that can shape population demography and abundance, and is an integral determinant of ecosystem functions (Engen and Stenseth 1989, Svanback and Persson 2004, Bolnick and Araujo 2011). Individuals exhibit differences in prey preference and prey acquisition efficiency, which, if heritable, may be a target of selection and ultimately promote ecological specialization (Fox and Morrow 1981, Smith and Skulason 1996, Devictor et al. 2010, Elmer et al. 2010, Machovsky-Capuska et al. 2016, Sexton et al. 2017). Large-scale disturbances in community structure, e.g., as a result of climate change (Sydeman et al. 2015) alter food web structures and the composition of available resources (Daufresne et al. 2009, Pershing et al. 2015, Bentley et al. 2017), forcing species to rapidly adapt to new diet landscapes. Therefore, understanding the underlying mechanisms shaping food acquisition strategies is fundamental to evolutionary biology and vital for predicting species survival in a changing world.

If heritable, the inter-individual variation in resource acquisition strategies may have complex evolutionary consequences mediated by trade-offs between energy gain and survival across density-and frequency-dependent fitness landscapes (Mousseau et al. 2000, Reznick and Ghalambor 2001, Reznick 2016, Sexton et al. 2017). For example, increased boldness to improve resource acquisition success may come at the expense of higher predation risk, the fitness costs of which may be linked to predator densities (Gotthard 2000, Carter et al. 2010, Bolnick et al. 2011). Likewise, the composition and abundance of available resources may alter the demographic structure of a population (e.g., Heino and Kaitala 1999, Enberg et al. 2012). For example, fast growth at an early age, e.g. as a result of abundant food sources during the initial stages of life, may result in early maturation and hence a younger age at reproduction. In contrast, resource limitation due to high population densities results in increased allocation to somatic growth to improve size-dependent intrapecific competition (e.g., Reznick and Endler 1982).

Ontogenetic diet shifts in organisms may be viewed as a special type of resource acquisition strategy in which diet variation is expressed as a function of age. Ontogenetic diet shift is a significant source of variation in species’ diet breadth, especially among size-and age-structured organisms, such as fishes. In general, relatively large and/or old individuals shift towards feeding at higher trophic levels and/or on larger prey items to maintain a positive energy balance (Werner and Gilliam 1984, Mittelbach and Persson 1998, Jensen et al. 2012). Under changing food-web dynamics, diet specialization among different age groups may substantially influence the demographic structure and life history diversity (Sanchez-Hernandez et al. 2019). For example, changes in resource composition that favour younger age groups would improve growth and subsequently increase the rate of maturation and the probability of survival at early ages. Ontogenetic diet shift is associated with a suite of changes in an individual’s morphology, physiology and behaviour to maximize the efficiency of particular resources at a given ontogenic stage, perhaps at the expense of reduced efficiency at other stages (Claessen and Dieckmann 2002).

If ontogenetic diet variation has a genetic basis, then some individuals in a population may be selected for high prey acquisition efficiency early in their life history (via physiological or morphological trade-offs towards efficient exploitation at earlier stages), even if this may come at a cost of compromised energy acquisition at later stages in life (Claessen and Dieckmann 2002). We predict that such genetically driven trade-offs in resource acquisition efficiency between early and late stages mediate the age structure (i.e. maturation timing) and abundance within and among populations and maintain genetic variation in resource acquisition strategies, but such examples in the wild are rare.

Atlantic salmon (*Salmo salar*) is a fish species recognized as a diet generalist and an opportunistic feeder with extensive ontogenetic and stage-and space-structured individual variation in diet breadth (Erkinaro et al. 1997, Jacobsen and Hansen 2001, Haugland et al. 2006, Hvidsten et al. 2009, Rikardsen and Dempson 2010, MacKenzie et al. 2012). At sea, where most growth occurs, salmon increasingly feed on prey at higher trophic levels as they grow and age (Jacobsen and Hansen 2001, Rikardsen and Dempson 2010). The time salmon spend at sea prior to maturation (sea age at maturity) also varies greatly within and among populations (Friedland and Haas 1996). Although the functional and physiological basis underlying age at maturity is not entirely known, it is considered to be a threshold trait, whereby higher lipid deposition is associated with early maturation (Friedland and Haas 1996, Jonsson et al. 1997, Thorpe et al. 1998, Taranger et al. 2010, Jonsson and Jonsson 2011). Therefore, variation in resource acquisition may be a strong determinant of the life history variation in salmon and a trait via which natural selection can act and result in adaptive genetic changes in populations.

In Atlantic salmon, two genomic regions on chromosomes 9 and 25 have been identified to have a disproportionate influence on life history strategy and population differentiation within and among populations (Ayllon et al. 2015, Barson et al. 2015, Czorlich et al. 2018, Pritchard et al. 2018, Aykanat et al. 2019). The so-called *vgll3* and *six6* genomic regions are named after the most prominent genes in their respective haploblocks on chromosomes 25 and 9, respectively. The *vgll3* genomic region on chromosome 25 has been shown to be associated with age at maturity (intially by GWAS, see: Ayllon et al. 2015, Barson et al. 2015), iteroparity (Aykanat et al. 2019), and precocious male maturation in Atlantic salmon (Lepais et al. 2017, Debes et al. 2019). This genomic region also exhibits strong spatial divergence (Barson et al. 2015, Pritchard et al. 2018), and it has recently been shown to have been affected by natural selection over the last 36 years (equivalent to 4-6 salmon generations) in parallel to the changing age structure in a large salmon population (Czorlich et al. 2018). The *six6* region on chromosome 9 is associated with sea age at maturity at the population level as a result of the strong correlation between the average allele frequency and average maturation age of populations (Barson et al. 2015). This region also exhibits the strongest signal of differentiation among European populations (Barson et al. 2015) and Tana/Teno River populations (Pritchard et al. 2018) and is hence distinguished as a critical genomic region for local adaptation. Genes found in these haploblocks appear to have a role in adipose or energy metabolism regulation in other organisms. The *vgll3* gene is an adipocyte inhibitor, the expression of which is correlated with body weight and gonadal adipose content in mice (Halperin et al. 2013). Recently, a strong selective sweep near the *vgll3* gene was postulated to be due to energy metabolism effects in humans in Mongolia (Nakayama et al. 2017). In turn, genes in the *six6* genomic region are involved in cell growth, cell differentiation, apoptosis in human cell lines (*PPM1A*, Lin et al. 2006), and myogenesis and skeletal muscle cell proliferation in zebrafish (*six1b*, Ridgeway and Skerjanc 2001, Bessarab et al. 2004, O’Brien et al. 2014) and act as an evolutionarily conserved regulator of eye development and the pituitary–hypothalamic axis (*six6*, Serikaku and Otousa 1994, Toy et al. 1998, Gallardo et al. 1999). Collectively, this suggests that the *vgll3* and *six6* haploblocks might have broad-scale roles in reproductive and life-history strategies in Atlantic salmon. However, how polymorphism in these regions may be translated to functional differences expressed in the wild is unclear.

Here, our objective was to test whether age-dependent differences in food acquisition efficiency are associated with the *vgll3* and *six6* genomic regions and discuss their role in explaining the genetic variation in age structure. We achieved this goal by assessing stomach content data from adult Atlantic salmon sampled along the coast during spawning migration and genotyping the same individuals for the *vgll3* and *six6* genomic regions using a targeted sequencing approach. Using a modelling framework that accounted for potentially confounding environmental and phenotypic variables, we tested whether variation in diet and resource acquisition strategies had a genetic component explained by the age at maturity-linked genomic regions. Elucidating the genetic interplay between age at maturity and diet breadth is crucial to better understand the dynamics and evolution of ecological specialization and to better predict future demographic changes in Atlantic salmon populations under climate change.

## Materials and Methods

### Sample collection

As part of a larger effort within the project “Sea salmon fishery, resource and potential (Kolarctic)”, Atlantic salmon (*Salmo salar*), on their return migration to spawning grounds, were sampled and stomachs were collected between mid-May and late July in 2008 by local sea fishers with bend nets or bag nets along the Finnmark coast, northern Norway (Svenning et al. 2019, Figure 1). Sampled fish were measured (fork length, cm) and weighed (g); their sex and maturity were identified, and stomachs were frozen for later diet analysis. In addition, scales were sampled from all fish for sea age determination, categorization as wild or farmed fish according to ICES guidelines (ICES 2011, Svenning et al. 2019), and genetic analysis. The species composition of the diet was then identified to species by visual inspection of the morphology of prey remains and otoliths which were compared to a reference collection with known species identity, with uncertain cases further inspected using guide books (Härkönen 1986, Pethon and Nyström 2005). All prey items, including unidentified digested remains were weighted. The identifiable portion of the diet in the dataset was overwhelmingly comprised of four fish species: sand eel (*Ammodytes* spp.), capelin (*Mallotus villosus*), herring (*Clupea harengus*), and haddock (*Melanogrammus aeglefinus*, see Results section for details). In the interest of analytical brevity, a few rare prey species were handled as follows: one gadoid fish was grouped with haddocks, both of which belong to the Gadiformes order, and negligible amounts of krill, other crustaceans, and Liparidae (0.2% of the total stomach weight) were categorized together with the unidentified material.

**Figure 1:**
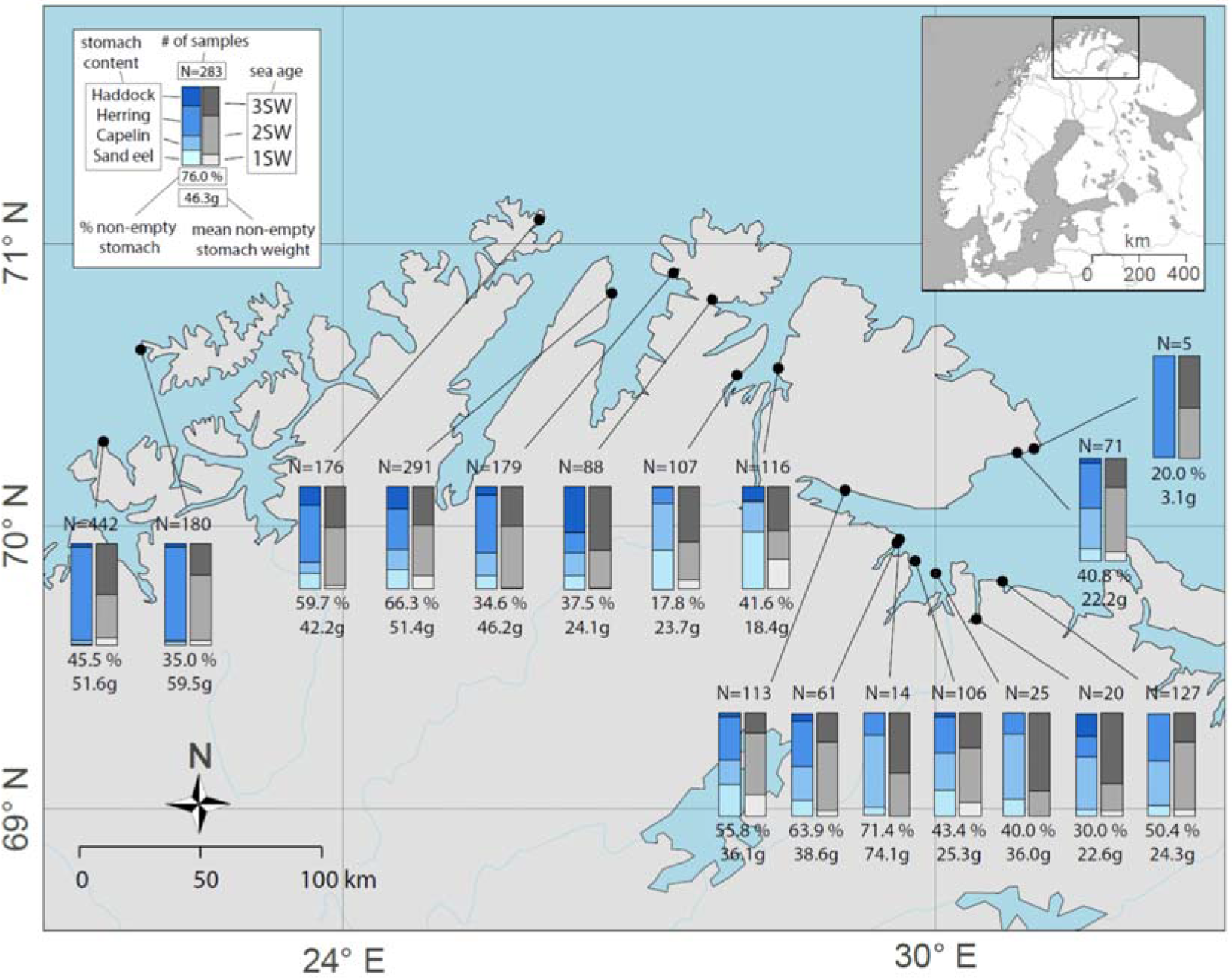
Map of the study region including sampling locations and a description of age and stomach content distributions and sample sizes in each sampling location. See the inset key for a description of the parameters of the spatial distributions given on the map.

### DNA extraction, microsatellite genotyping, and SNP genotyping by targeted sequencing

DNA was extracted from scales either using a QIAamp 96 DNA QIAcube HT Kit (Qiagen) following the manufacturer’s protocol or according to (Elphinstone et al. 2003). Microsatellite genotyping of 31 markers was performed as outlined in (Ozerov et al. 2017). Samples were further genotyped by targeted sequencing at 173 SNP markers and the sex determination locus (sdy) using a GTSeq approach (Campbell et al. 2015) as outlined in (Aykanat et al. 2016), with some modifications so the genotyping panel was compatible with the Illumina platform. More specifically, 174 genomic regions were first amplified in one multiplex PCR using locus-specific primers with truncated Illumina adapter sequences and using primer concentrations re-optimized for the Illumina platform (Supp. Table 1). The PCR products were then treated with Exonuclease I and FastAP Thermosensitive Alkaline Phosphatase (Thermo Fisher) to remove unused primers and nucleotides. After the treatment, the products were re-amplified with adapter-specific primers containing Illumina and sample-specific dual-indexes. The index set was optimized using the BARCOSEL software (Somervuo et al. 2018). The PCR products were then pooled, purified and quantified with a Qubit 2.0 fluorimeter (Thermo Fisher) and analysed on a fragment analyser (Agilent Technologies). The pooled library was then size selected using BluePippin (Sage Sciences) to remove short unspecific products and checked on a fragment analyser. Finally, samples were single-end sequenced using a 150-cycle high-output sequencing kit on a NextSeq 500 Illumina Sequencer following the manufacturer’s guidelines. Loci with coverage over 12x were scored as in (Aykanat et al. 2016). To calculate coverage for each SNP, raw genotype files (fastq) were scanned for every SNP, and the coverage was determined by counting sequences that matched SNP’s forward and reverse primer sequences, and the 9 bp region around the SNP site. Finally, genotypes were scored based on coverage ratios between alleles: Ratio_Cov(allele1/allele2)_ >10 was assigned as homozygous for allele 1, Ratio_Cov(allele1/allele2)_ <0·1 was assigned as homozygous for allele 2, Ratio_Cov(allele1/allele2)_ between 0·2 and 5 was assigned as heterozygous and any proportion in between was discarded (see also, Campbell et al. 2015).

Two focal SNPs used in the analyses were *vgll3*_TOP_, which exhibits the strongest signal of association with age at maturity in the *vgll3* genomic region, and *six6*_TOP.LD_ in the *six6* haploblock on chromosome 9, the region that exhibited the second strongest association with sea age at maturity prior to population structure correction and is 34.5 kb away from and in complete linkage disequilibrium with the *six6*_TOP_ SNP reported in Barson et al. (2015).

### Genetic stock identification (GSI)

In total, 2023 samples that had greater than 80% success in regard to microsatellite genotyping were assigned to their population of origin with 31 microsatellite markers as described in (Svenning et al. 2019) using the Bayesian GSI methodology described in (Pella and Masuda 2001) and implemented in cBayes 5.0.1 (Neaves et al. 2005). In brief, the samples were allocated into 18 analysis groups, that is, the combination of two time periods (May-June and July) and 10 fisheries regions, with each group consisting of 30 to 288 samples for analysis. The GSI analyses were performed using five independent Monte Carlo Markov chains of 100K iterations starting from three random stocks (StartStock parameter), and the last 10K iterations of each chain were combined and used to assign individuals to their population of origin to remove the influence of initial starting values. The baseline population data for the GSI analysis included genetic information on 185 Atlantic salmon populations spanning from the Pechora River (Russia) in the east to the Beiarelva River (Norway) in the west (see details in Ozerov et al. 2017).

The probability (p) threshold for assignment of an individual to a population was set at ≥0.7 following Vähä et al. (2011, 2014) and Bradbury et al. (2015). The 30% of individuals assigned to a population with lower confidence were kept in the dataset with the highest ranked population assigned as the population of origin. Samples with no assignment due to low genotyping success with microsatellites (4%) were assigned a population of origin using genotype information from the SNP panel. In such cases, individuals were assigned to the population in which they exhibited the highest genetic similarity. This was measured according to the average genetic similarity of focal individuals to the individuals in each population (as inferred with the GSI analysis in the previous step) using the *A.mat* function in the rrBLUP package (Endelman 2011) A small subset individuals (N=16, < 1%) with poor genotyping success with SNPs (less than 50 SNPs with high quality genotypes) was randomly assigned to a population, in which population assignment probability is weighted over the total number of individuals that were assigned by GSI. The effect of including incomplete population assignment was assessed for the main analysis by repeating the analysis but only including confidently assigned individuals.

Missing data points for some variables were inferred from highly correlated variables. In that regard, missing sea age information (i.e., due to unclear formation of sea annulus for detecting the correct sea-age for some first time spawners) was inferred from length data for 15 (0.7%) individuals, where the likelihood of age, given the length, was substantially higher (>20 times) for the inferred age group than for other age groups. Additionally, for 21 (1.0%) individuals with missing length data, fit using coefficients of log(weight) to log(length) regression (adjusted *R*^2^ = 0.94) was used to estimate the length information from the weight data. Finally, data for 33 (1.6%) individuals with missing *vgll3_TOP_* genotype scores were inferred from the genotype score of an adjacent SNP marker in the genotyping panel, vgll3_Met54Thr_, which is in close physical proximity to *vgll3_TOP_* with high linkage disequilibrium (*r*^2^ = 0.79).

### Genetic and ecological basis of the diet scope

Unless otherwise noted, all statistical analyses were performed in R software v.3.2.5 (R Core Team, 2018). Either a two-component hurdle model (with binomial and the conditional negative binomial components) using the *glmmTMB* package (Brooks et al. 2017) or a binomial model (with a log link function) using the *gamm* function in the *mgcv* package (Wood 2011) was employed as the statistical model. In all models, population of origin was included as a random term to account for background population effects. To control for spatio-temporal variation, sampling location (longitude) and the day of sampling (Julian day, zero centred, and scaled to one standard deviation) were included as smoother terms. Longitude, which explained 90.7% of the spatial variation (i.e., sampling locations mostly occurred along a longitudinal axis (see Figure 1) and were included in the models as a surrogate for the two-dimensional spatial distribution to decrease the parametrization of the model. In addition to including the genetic variation in the *six6* and *vgll3* genomic regions additively in the model (i.e., genotypes coded as a continuous factor with heterozygotes coded as the average of two homozygotes), age at maturity (e.g. Fleming 1996) and residual length (log transformed total length after controlling for age at maturity) were also included in the model as categorical and continuous variables, respectively. All numeric variables were centred and scaled. For both genomic regions (*six6* and *vgll3*), alleles associated with late and early age at maturity were labelled as *L* and *E*, respectively.

The general model structure was as follows:

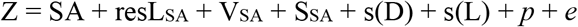

where Z is the vector of response variables given as a function of sea age (SA), scaled residual length and *vgll3* and *six6* genotypes nested within sea age (resL_SA_, V_SA_ and S_SA_, respectively), the smoother functions of sampling day (D) and location (L), and normally distributed random variance due to the population (*p*) and individual (residual) effects. The genotypes are coded additively as 1, 2, and 3 (for EE, *EL* and *LL*, respectively). In the model, the scaled genotypes and residual length were analysed independently within sea age group (i.e., nested model), which provided a statistical framework suitable for testing hypotheses related to ontogenetic diet structure. In this model, a small number of 4SW individuals (N = 15, 0.7%, SW denotes number of winters salmon spent at sea prior to sampling.) was grouped and analysed within the 3SW for statistical coherence of the nested model. Models also accounted for spatio-temporal variation in the diet with smoothing spline functions. When using the *gamm* package, which provides a platform for generalized additive models, days and location were modelled with a smooth function (*s()*). When using *glmmTMB* package, which provides a platform for hurdle models but cannot directly model the smoother functions, an orthogonal spline design matrix with a low-rank thin-plate function was generated using the *spl* function in the *MCMCglmm* package (Hadfield 2010) in R and included in the model as fixed terms as a surrogate for the spatio-temporal spline functions. The number of knot points (*k*, which defines the curvature of the spline function) was set to five for both variables, but the results were robust to an increase in the *k* value, which did not qualitatively change the results (data not shown).

A number of variables pertaining to the diet content data were used as response variables in this study. Conceptually, these variables are linked to different aspects of diet acquisition mechanisms of this species (e.g., Arrington et al. 2002) and may indicate different functional aspects associated with performance and life history variation among individuals.

#### 1) The presence and amount of total diet content in the stomach

We used a two-component hurdle model to simultaneously account for stomach content quantity and empty stomach probability. By this, we tested for the prevalence of the “feast and famine” diet acquisition strategy in Atlantic salmon as a function of ontogeny and genotype, whereby large piscivorous fish species are predicted to experience prolonged periods with empty stomachs in the interest of acquiring a high quantity of food (Arrington et al. 2002, Armstrong and Schindler 2011). Both components in the hurdle model included the same set of covariates (as described above). A logistic model with a log link was used to model the probability of the presence of a prey item, whereas a zero truncated negative binomial model with a log link was utilized as the conditional component. In this analysis, the total stomach weight, which had excess zero elements and a right-tailed continuous distribution (ranging from 0.5 to 393.3 g), was transformed to a discrete distribution by arbitrarily binning the total weight in 10 g increments, with zero stomach content set as the first bin at a value of zero (Supp. Figure 1). This transformation provided a distribution that can be modelled with the hurdle framework in the *glmmTMB* package (Brooks et al. 2017). Finally, we also repeated the analysis by only including confidently assigned indiviuals in order to assess the robustness of model to incomplete population assignment.

#### 2) Total number and average weight of prey items in the stomach

An increase in the total prey weight in the stomach can be explained either by an increase in prey number or an increase in the average prey weight. Therefore, we next investigated the contribution of these two components in terms of explaining the model using the same statistical framework as above. Similar to the total prey weight, the average prey item weight (ranging between 0.2 and 300.2 g) was also transformed to discrete units by arbitrarily binning the data at 5 g intervals, with zero stomach content set as the first bin (Supp. Figure 1).

#### 3) Relative prey composition

Finally, to test whether sea age, size at age, or genotype is associated with specific prey species, we modelled the prey composition, measured as the proportion of a specific prey species contributing to the stomach content weight. The proportion of each of the four prey species in the total prey weight was modelled as a response variable using binomial regression in the mgcv package (Wood 2011)

Extensively digested, unidentified content in the stomach (4.5% of the total stomach weight) was not treated as diet material in order to accurately reflect the recent feeding activity (e.g., Jacobsen and Hansen 2001). For all models, the effect size and confidence intervals were calculated with 10,000 parametric permutations of the model coefficients. To account for potential spurious inflation associated with genotype, i.e. due to cryptic family structure, the analytical pathway was repeated using 168 independent and putatively neutral markers that are present in the SNP panel, and focal SNPs were ranked across the background genetic effect (by comparing the genetic model and the null model at each SNP marker).

## Results

The final dataset contained 2121 individuals after excluding previously spawned and escaped farmed salmon. In the final dataset, 93.3% of the samples had visibly detectable developing gonads, confirming concordance between sea age and sea age at maturity. A total of 1372 (64.7%) individuals were confidently assigned to a population of origin (p > 0.7), and (N=651) Individuals assigned to a population with lower confidence (p < 0.7, 30.7%). A further 82 individuals (3.9%) with low genotyping success with microsatellites were assigned a population of origin using SNP data and 16 individuals (< 1%) lacking reliable genotypes with either set of genetic markers were randomly assigned to a population (see Materials and Methods for details).

### Stomach content analysis

Out of the 2121 individuals examined in the final dataset, 992 individuals had identifiable prey items in their stomachs (46.8%). Four fish species, sand eel (*Ammodytes* spp.), capelin (*Mallotus villosus*), herring (*Clupea harengus*), and haddock (*Melanogrammus aeglefinus*), comprised the bulk of the diet content, representing 42.2 kg of the 44.2 kg quantified diet content (95.5%). In total, there were 2843 identifiable prey items in the datasets, with sand eel being the most abundant and herring being the largest percentage by weight (Supp. Figure 2). On average, prey weight significantly differed among species, with haddock being the heaviest, followed by herring, capelin, and sand eel (Supp. Figure 2).

### Prey probability and weight in the stomach as a function of sea age, size at age, and genetic variation

The two-component hurdle model revealed a striking negative relationship between the probability of non-empty stomach (e.g. presence of identifiable prey item in the stomach) and prey weight (g) in the stomachs of Atlantic salmon as a function of sea age (Figure 2a-b, Supp. Table 2). Young age groups were more likely than older age groups to have prey in their stomach. 1SW individuals were 1.45 (1.07 - 1.95, 90% CI, *p* = 0.020) and 2.39 (1.73 - 3.30, 90% CI, *p* < 0.001) times more likely to have any prey item in their stomachs than 2SW and 3SW individuals, respectively, and 2SW individuals were 1.65 times more likely to have a prey item in their stomachs than 3SW fish (1.35 - 2.00, 90% CI, *p* < 0.001). The decrease in non-empty stomach was significantly associated with residual size variation within the 2SW age group (*p* = 0.027, Figure 2e), with larger individuals having empty stomachs more often than the smaller-sized fish in the same age group.

The conditional truncated negative binomial model suggested that young age groups had significantly less prey in their stomachs than older age groups despite a higher likelihood of having non-empty stomachs (Figure 2b). The contrasting results between the zero-inflated and truncated negative binomial components suggested that resource acquisition strategies differed among age groups. The model estimated, on average, 9.9 g (7.0 – 13.9, 90% CI), 24.8 g (21.8 – 28.6, 90% CI), and 41.5 g (33.8 – 50.9, 90% CI) of prey items in the stomachs of 1SW, 2SW and 3SW fish, respectively, all of which were highly significantly different from one another (p < 0.001). Residual length at age also appeared to be a predictor of prey content, but only significantly so in the 2SW age group (*p* = 0.036, Figure 2b).

**Figure 2:**
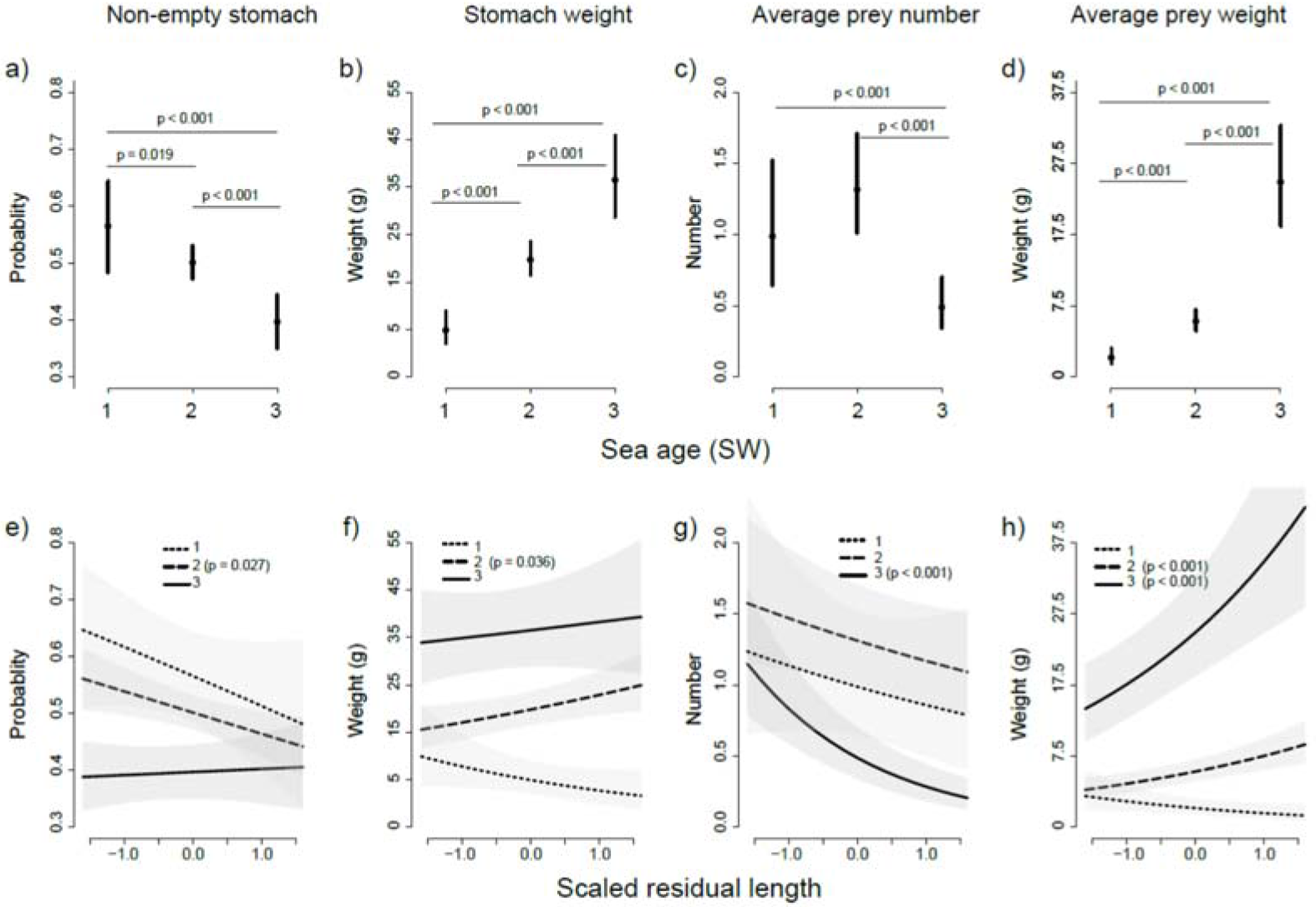
Changes in four stomach content variables in relation to sea age (a) and size at age (b) and associated 90% confidence intervals. Only p-values < 0.1 are given in the figures. Binned prey weights are given after back transformation to the original scale (i.e., grams). Note that estimates in the last three columns (stomach weight, average prey number and weight) are for non-empty stomachs only.

The *six6*\**L* allele, which has a higher frequency in populations with an older sea age at maturity (Barson et al. 2015), was associated with an increase in the probability of non-empty stomach in an age-dependent order, with a more pronounced effect in younger age groups (Figure 3a). Allelic substitution from *E* to *L* in the *six6* genomic region (i.e. change in effect size by changing one allele of the genotype) increased the probability of prey occurring in the stomach by 1.56 (1.06 – 2.30, 90% CI, *p* = 0.057) and 1.25 times (1.08 - 1.45, 90% CI, *p* < 0.014) in the 1SW and 2SW groups, respectively (Figure 3a). Both age groups exhibited significant or near significant age-dependent genotype effects relative to the 3SW group (*p* = 0.036 and 0.051, respectively, Figure 3b). Strikingly, the probability of prey in the stomach in relation to size at age (Figure 2e) was in the opposite direction to the *six6*\**L* effect (Figure 3a) despite the two (*six6* and size at age) exhibiting a significant positive correlation (Supp. Table 3), suggesting the occurrence of complex, contrasting effects of *six6* genetic variation across different phenotypic classes.

**Figure 3:**
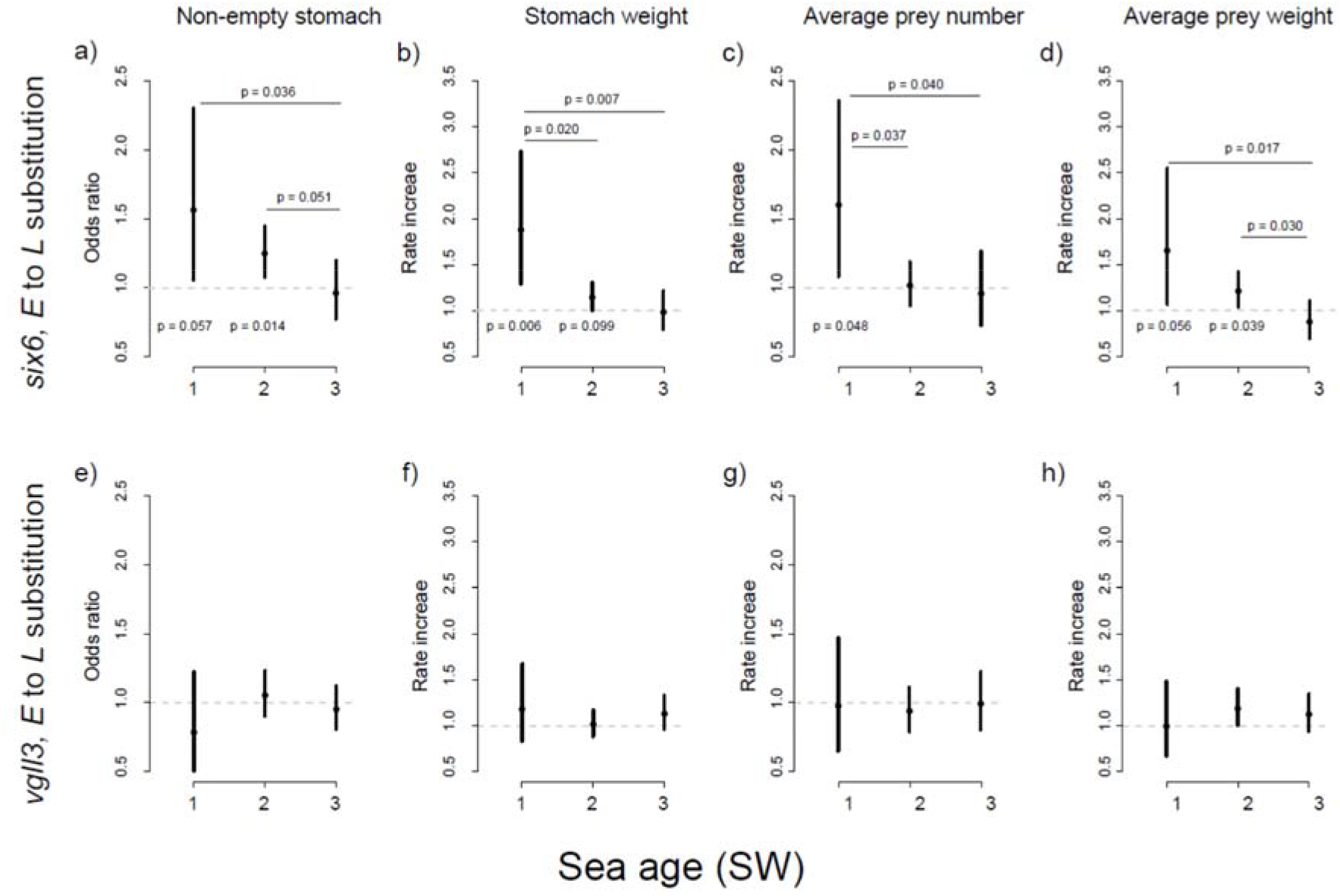
Changes in four stomach content variables for differing sea age classes in relation to allelic substitution in *six6* (a) and *vgll3* (b) genomic regions and associated 90% confidence intervals. Only p-values < 0.1 are given in the figures. Note that estimates in the last three columns (stomach weight, average prey number and weight) are for non-empty stomachs only. P-values above the plots indicates comparative difference in genetic effects between sea age classes, and below indicates the significant genetic effect within age classes (no odds change in relation to allelic substitution). Same odds (odds ratio = 1) indicated by a dashed line.

The conditional model suggested that the *six6*\**L* allele was also associated with increased total stomach weight in the young age groups (1SW and 2SW) but not in the 3SW group (Figure 3b). The allelic substitution effect from *E* to *L* was significant and associated with a 1.87-fold (1.29-2.73, 90% CI, *p* = 0.006) increase in prey weight in the 1SW group and was marginal in the 2SW group, associated with a 1.14 (1.00-1.30, 90% CI, *p* = 0.099) increase in prey weight (Figure 3b).

The *vgll3* genomic region was not associated with diet content variation, suggesting no causal link between the two (Figure 3e-h). However, selection on diet may still exert evolutionary change in the *vgll3* genomic region, via correlated response to selection (Lande and Arnold 1983), as a result of phenotypic covariation between diet and age at maturity and length at age. Accordingly, the effect of *vgll3* was significant when these co-varying phenotypes were not accounted for in the model (Supp. Table 4). Spatio-temporal variance in the dataset was substantial in explaining diet variation in both components of the hurdle model (Supp. Table 2, see also Supp. Figure 3). In general, diet presence and quantity were the highest at the westerly end of the distribution, with a gradual decrease towards the east. At the temporal scale, sampling days in the middle of the sampling period were associated with a higher presence and quantity of diet in the stomach (Supp. Figure 3).

Population of origin was not a significant source of diet variation and explained only a fraction of the total variation in diet content (Supp. Table 2, ΔAIC = 3.91, LRT_2,2057_ = 0.09, p = 0.96). When the analysis was performed with samples assigned to a population of origin with high confidence (N = 1372), the variance due to population was similarly small (see Supp. Table 5 for total stomach weight as the response variable) and the model was also less parsimonious than a model without the population effect, as assessed by comparing the model fit by difference in their Akaike information criterion and likelihood ratio test (ΔAIC = 3.42, LRT_2,1340_ = 0.58, p = 0.75). Likewise, the relation between total stomach weight and *six6* genetic variation was relatively robust for the full dataset (N=2057, Supp. table 2) and dataset only including individuals with high population assignment confidence (N=1372, Supp. table 5).

In our framework, digested, unidentified material in the stomach was not included in the analysis (e.g., Jacobsen and Hansen 2001). However, the results were qualitatively similar when digested material was included in the analysis (Supp. Table 6). A model including sex was less parsimonious and the term was not included as a parameter in the model (ΔAIC = 0.75). Finally, when the fit of the genetic models (*six6* and *vgll3*) was compared to the putatively neutral SNPs in the panel, *six6* ranked first out 167, confirming its significance, while *vgll3* was only ranked 123^rd^ (Supp. Figure 4).

In general, both the number of prey items and the increase in the individual prey weight contributed to the variation in the total stomach weight (Figures 2c-d & 3c-d, Supp. Tables 7 & 8). The 3SW age group was associated with significantly fewer prey items (0.49 prey items, 0.34-0.71, 90% CI) than the 1SW (0.99 prey items, 0.65-1.52, 90% CI, *p* = 0.004) and 2SW age groups (1.32 prey items, 1.01-1.71, 90% CI, *p* < 0.001), but the average prey weight was significantly heavier (27.40 g, 21.24-35.35, 90% CI) than that in the 1SW (2.68 g, 1.79-4.06, 90% CI, *p* < 0.001) and 2SW (7.83 g, 6.45-9.53, 90% CI, *p* < 0.001) salmon. The average prey weight, but not the prey number, was also significantly different between the 2SW and 1SW age groups (*p* < 0.001, Figure 2c-d). Size within age group also significantly influenced the number and size of prey. Larger fish within the 3SW age group had fewer (*p* < 0.001) but heavier (*p* < 0.001) prey items in the diet than smaller fish, and larger 2SW individuals consumed smaller prey items (*p* < 0.001, Figure 2g-h, Supp. Tables 7 & 8).

The number and size of prey items was also explained by the *six6* genotype in an age-dependent manner, with a more pronounced effect in the relatively young age groups. The *E* to *L* substitution in *six6* was associated with a 1.60-fold (1.08-2.36, 90% CI, *p* = 0.048) increase in prey number in the 1SW age group (Figure 3c). The allelic substitution was also associated with 1.65-fold (1.07-2.56, 90% CI, *p* = 0.056) and 1.22-fold (1.04-1.42, 90% CI, *p* = 0.040) increases in average prey weight in the 1SW and 2SW age groups, respectively, which was significant compared to that observed in the 3SW age group (*p* = 0.018 and 0.030, respectively, Figure 3d). Genetic variation in *vgll3* was not significantly associated with average individual prey weight or prey number after controlling for age at maturity (Figure 3e-h).

### Relative prey composition as a function of sea age, size, and genetic variation

Prey composition varied substantially across different age groups, suggesting a change in prey composition as the fish grow older (Figure 4). In general, older age groups were more likely to prey on herring and haddock (Figure 4a-b), while younger age groups preyed on capelin and sand eel (Figure 4c-d). The same pattern was observed within age groups, (e.g. larger fish within an age group had proportionally more herring and haddock than smaller fish in the same age group) albeit generally not significantly (Supp. Figure 5), suggesting that size may be a contributing factor explaining prey composition. In all analyses, spatio-temporal variation was a significant component explaining the prey composition (Supp. Table 9).

**Figure 4:**
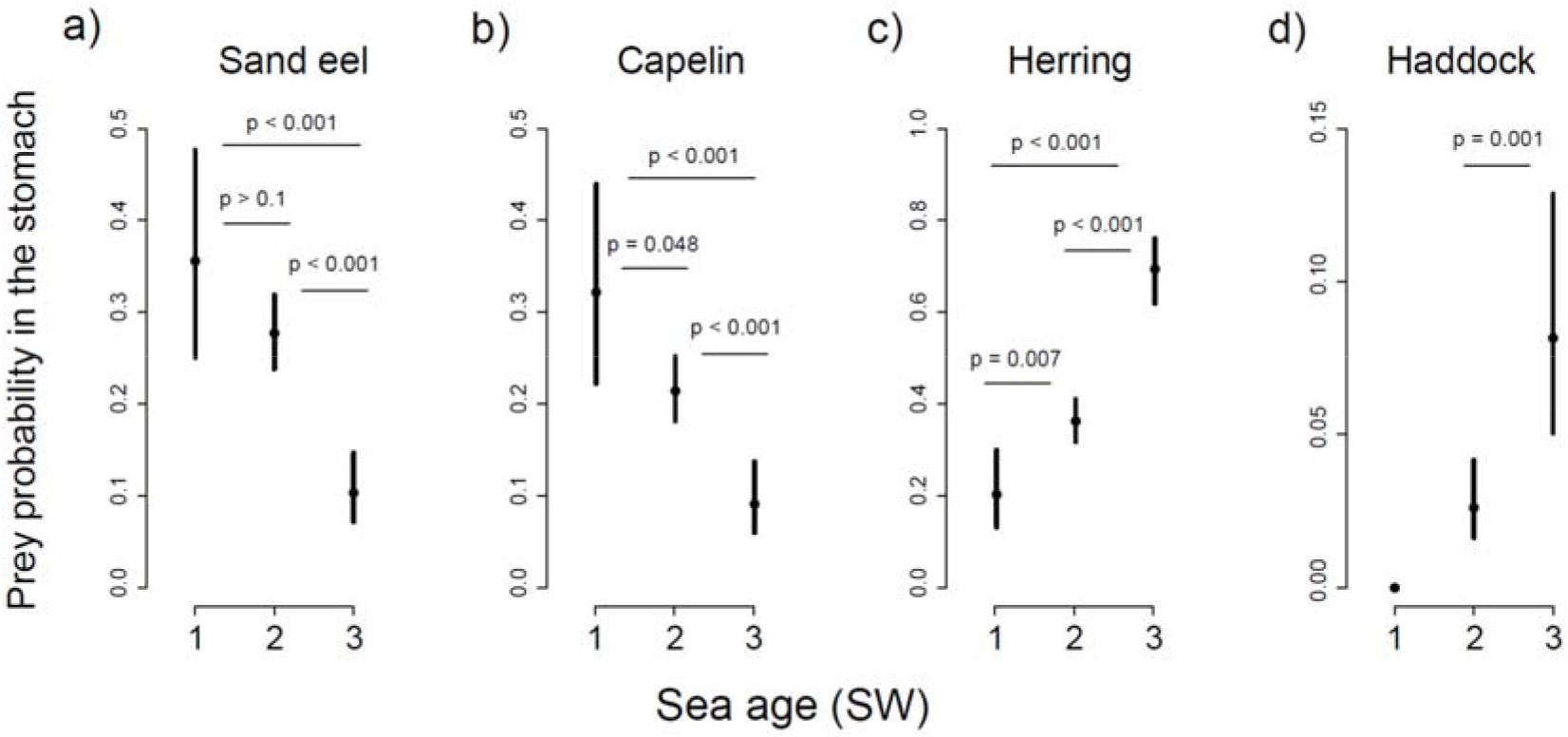
Relative prey composition for the four key prey species, measured as the proportion of each prey species contributing to the stomach content weight, in relation to age at maturity. Note that estimates include only non-empty stomachs.

Genetic variation in *six6* and *vgll3* did not appear to be a strong predictor of prey composition, but some notable associational trends existed for the two genomic regions (Supp. Figure 6, Supp. Table 9). Particularly, *E* to *L* substitution in *vgll3* is associated with 1.31 times (1.01-1.70, 90% CI, *p* = 0.089) and 1.49 times (1.03-2.16, 90% CI, *p* = 0.076) fewer capelin in the stomach relative to other prey species in the 2SW and 3SW age groups, respectively (Supp. Figure 6). Additionally, there was a significant age-dependent preference for capelin over herring associated with the *E* to *L* substitution in the *six6* genomic region (Supp. Figure 6). When compared to putatively neutral SNPs in the genotyping panel, capelin composition modelled with *vgll3* ranked 12^th^ out of 164 SNPs (0.073, Supp. Figure 7), a value that is consistent with the analytically inferred p-value. Finally, when sea age and size at age were not controlled for, as expected, genetic variation in both the *vgll3* and *six6* genomic regions explained a substantial portion of the variation also in the relative prey composition (Supp. Table 10). This suggest the phenotypic covariance between and diet composition and age at maturity may exert a correlated response to selection at the life history genomic region (i.e. *vgll3*), despite not being causally linked to diet (i.e. Lande and Arnold 1983).

## Discussion

Quantifying resource acquisition via stomach content analysis is an integral component of trophic studies in ecology and evolution. In this study, we used stomach content information from a single time point as a proxy for diet acquisition. The analysis of diet content may be difficult for a number of reasons, such as the challenges in accounting for the diverse nature of diet content data, high percentage of digested items, and the diversity of metrics for statistical analysis (e.g., Rice 1988, Cortés 1997, de Crespin de Billy 2000, Baker et al. 2014). On the other hand, the relatively small number of diet species in our dataset, all fishes, and the small proportion of undigested material provided us with a robust quantification of the diet and a powerful statistical framework with ecologically relevant response variables. The three most common prey species in our dataset (e.g., herring, capelin and sand eel) were from similar trophic levels independent of size or ontogeny (Dommasnes et al. 2001, Bentley et al. 2017). In turn stable isotopes, which are sensitive to quantify diet variation across trophic levels, would have been unable to tease the observed variation apart. Although stable isotopes may effectively quantify diet variation at other life stages, or shed light on long-term dietary patterns, stomach content analysis is a more robust approach when diet composition is variable when considering similar trophic levels.

Our analyses indicate that diet acquisition strategies in the sea vary with sea age in Atlantic salmon and that this variation is associated with genetic variation in key life history genomic regions, particularly in the *six6* genomic region. The variation in diet explained by sea age and size at age was mostly concordant, suggesting that size is the major driver of diet variation, influencing both the quantity and species composition of prey (Figures 2 & 4). Atlantic salmon prey on heavier but fewer prey as they grow older and larger, which seems to be a strategy that comes at the expense of a reduced prey acquisition probability (Figure 2a-d, Huey et al. 2001, Arrington et al. 2002). This pattern is consistent with the so-called “feast and famine” strategy observed among large piscivorous fish species (Arrington et al. 2002). The feast and famine feeding strategy is suggested to be an adaptation to maintain a positive energy balance at a large body size, especially when the acquisition of energy-rich food sources is unpredictable (Armstrong and Schindler 2011). Large Atlantic salmon appear to adopt this strategy, which is likely beneficial in terms of balancing the increase in energy costs associated with a large body size. A suite of physiological adaptations and metabolic adjustments, such as increased digestion capacity (Armstrong and Schindler 2011) and fat storage (Bustard 1967), may be associated with this strategy (Wang et al. 2006). For example, it has been shown that piscivorous species that adopt a feast and famine strategy maintain a large digestive tract, which provides quick food utilization when abundant prey are encountered (Armstrong and Schindler 2011). This physiological trade-off seems to be evolutionarily favourable for large fish when the prey distribution is stochastic despite the energetic costs of sustaining excess and energetically expensive digestive tissue (Armstrong and Schindler 2011). The feast and famine strategy in large Atlantic salmon may also be facilitated by other mechanistic processes, such as the trade-off of a lower success rate linked to larger prey or a lower attack rate associated with increasing size. Nonetheless, variation in foraging strategies among different age groups results in a large diet breadth, efficient resource partitioning, and reduced intraspecific competition among age groups, which subsequently promotes their co-existence (e.g., Polis 1984, Smith and Skulason 1996, Svanback and Bolnick 2007). It is unclear what physiological or behavioural modifications are associated with the differential feeding strategies among the various age groups in Atlantic salmon. Nonetheless, changes in marine food webs may alter the density and composition of prey available to different age groups and hence alter the age-dependent selection landscape, potentially leading to adaptive changes in age structure. Our results confirm the value of Atlantic salmon as a model species to study the evolutionary physiology of starvation and feeding in response to environmental changes in the wild.

Salmon are generally considered to be diet generalists. However, the association between the life history genomic region *six6* and diet acquisition (as well as the suggestive acquisition with capelin prey preference and *vgll3*, see, Supp. Table 6) indicates that genetically controlled intraspecific diet specialization occurs in Atlantic salmon. This genetic variation may be linked to specialized dietary adaptations (e.g., physiological, morphological or behavioural) allowing the efficient utilization of diverse diet sources, resulting in increased niche breadth and reduced intraspecific competition (Dalmo et al. 1997, Bolnick et al. 2003, Bolnick and Fitzpatrick 2007, Svanback and Bolnick 2007). In particular, the *six6*\**L* allele was associated with increased content in the stomach, especially in young age groups, which was explained by both an increase in the number of prey items and an increase in the average prey weight in the stomach. Intriguingly, however, the *six6*\**L* allele was associated with a larger fish size within all sea ages (Supp. Table 3, see also, Barson et al. 2015). Given that larger size at a given sea age is indicative of higher performance and fitness in adult salmon (e.g., Fleming 1996; Mobley et al. 2020), the *six6*\**L* allele (being linked to larger size) may be expected to have a selective advantage over the *E* allele and is hence predicted to prevail within and among populations as a result of directional selection. However, the *six6* genomic region is highly variable within and among populations (Barson et al. 2015, Pritchard et al. 2018), and balancing selection appears to be the pervasive mode of evolution, with both alleles exerting fitness advantage with different life-history strategies (Barson et al. 2015). The results of this study suggest that the genetic variation in *six6* locus could be explained by antagonistic pleiotropy along the life cycle of an individual (i.e., fitness trade-offs within the lifetime of an individual associated with the *six6* region) or fluctuating selection across generations. For example, genetic variation in *six6* may be under balancing selection via antagonistic genetic correlations in diet acquisition over the life cycle. Indeed, some results indicate that individuals having high freshwater growth may show poor seawater growth (Einum et al. 2002), prescribing further hypothesis linking it to genetic variation in *six6* genomic region.

Although our results suggest superior performance of the *six6*\**L* allele at the adult stage, the opposite may be true at earlier life stages, when the prey species composition is substantially different, with significantly more invertebrates in the diet (Jacobsen and Hansen 2001, Rikardsen et al. 2004, Haugland et al. 2006), and different ecological drivers affect performance (Mittelbach et al. 2014, Sanchez-Hernandez et al. 2019). For example, if increased metabolic costs are linked with increased prey content in the stomach, a trait associated with *six6*\**L*, this may not be an optimal acquisition strategy in younger years (i.e. juveniles prior to, or in the early phases of marine migration) when the energy density of available prey cannot compensate for the greater effort (McNamara and Houston 1996, Enberg et al. 2012). Such antagonistic genetic correlations between early and late life histories (i.e. genotype-environment interactions) in diet acquisition may maintain polymorphism in the region. Alternatively, year-to-year variation in population-specific (e.g., density dependent) or ecosystem-level processes (i.e., prey composition, food-web dynamics) may alter the adaptive landscape of diet acquisition (Smith and Skulason 1996), resulting in fluctuating selection and a change in the direction of selection among alleles, which would maintain the genetic variation in the region. Indeed, in the year 2008, returning salmon had a notably older sea age at maturity in northern Norway than that observed in more recent years (i.e., the oldest in the last 20 years, see (Anon 2014). This is consistent with observed patterns that *six6*\**L* is linked with larger size, perhaps as a result of providing a selective advantage for that particular year class. However, more research is required to explicitly test this possibility. The *six6* genomic region is highly spatially differentiated among populations (Barson et al. 2015, Pritchard et al. 2018). This suggests that any selection acting on *six6* as a result of selection on diet acquisition would influence the fitness of populations differentially, correlated with their average allele frequency. Hence, genetic variation in *six6* may be linked to the differential survival of populations at sea and should be closely monitored in population management.

The genetic variation in the v*gll3* genomic region had no clear effect on diet quantity overall, but there was a marginal (p = 0.073), albeit consistent, association with higher capelin content in the stomach (Supp. Figures 6 & 7). Capelin is a key component in the Barents Sea ecosystem with critical bottom-up effect (Gjøsæter et al. 2016), and both salmon post-smolts and returning adults utilize them as a food resource (Rikardsen and Dempson 2010, this study) when they migrate through the coastal regions (Gjøsæter et al. 2016). Individuals with the *vgll3*L* allele, which is associated with a later sea-age at maturity (Barson et al. 2015), were marginally less likely to feed on capelin, particularly in older age groups (2SW and 3SW). In addition, in contrast to herring, which was the most common prey item in the stomachs of the individuals in the 2SW and 3SW age groups (Figure 4c), capelin is likely a lower-energy prey item for Atlantic salmon (Elliott and Gaston 2008, Hedeholm et al. 2011, Renkawitz et al. 2015). This supports the notion that *vgll3*L* may contribute to foraging adaptations to support the high-energy demands associated with the larger body size of late maturing fish. However, the mechanisms driving such a compositional difference have yet to be determined.

Overall, genetic variation in both life history genomic regions appears to have a role in intraspecific diet specialization, but the mechanisms remain to be clarified. Although the underlying mechanism of diet specialization is complex (e.g., Mittelbach et al. 2014) and challenging to disentangle, performance trade-offs across different ecological settings and the life cycle are likely driving the life-history variation associated with these genomic regions. Analyses of datasets collected in the wild may suffer from confounding effects that co-vary with both the response variable and parameters of interest and generate spurious associations if they are not accounted for. In this study, a highly controlled linear model was employed to account for environmental and intrinsic parameters with potential confounding effects. First, resource availability at sea, e.g., the prey species density and distribution, is highly variable across time and space, even at small scales. Likewise, during their return migration, Atlantic salmon may be non-randomly distributed in relation to their life history, genotype, and population of origin (Svenning et al. 2019). For example, relatively early run timing is linked to both later maturation (Jonsson and Jonsson 2011) and *six6*\**L* (Cauwelier et al. 2018, Pritchard et al. 2018), a pattern that concordantly holds in our dataset (Supp. Table 11). Hence, the analytical framework controlled for spatio-temporal variation and also accounted for non-linear changes through space and time (Supp. Figure 3). Similarly, sea age at maturity and size within an age group exhibited strong links with both diet and genetic variation in the life history genomic regions. By accounting for these phenotypes in the model, we were able to exclude the possibility of modelling the genetic variation via the effects of these intermediate phenotypes. Therefore, our framework was rather robust to drawbacks related to confounding factors observed in wild settings. Finally, variation in diet due to the population of origin was also accounted for in the model as a random intercept but did not explain significant diet variation at sea (Supp. Tables 2 & 5).

On the other hand, the significance between diet variation and life history genomic regions were often only marginally significant. Therefore, further research is warranted to validate these results. Further, genetic variation outside these two major maturation-timing loci and potential pleiotropic effects remain unexplained, which could be explored with a genome wide approach in future studies. Likewise, similar set-ups at different life history stages, under different feeding regimes, or in common garden conditions, would help to elucidate the effect of genotype-environment interactions that may have been overlooked in this dataset.

Marine ecosystems, which are composed of mostly poikilothermic species, are sensitive and highly responsive to temperature-driven changes (e.g., Clarke 2003, Sydeman et al. 2015). The Arctic region is particularly sensitive to global climate change (Polyakov et al. 2010), with significant anthropogenic effects further shaping the marine food webs in the region. Over the last 40 years, the abundance of Atlantic salmon has been declining, and the age structure has been shifting towards a younger age at maturity (Chaput 2012, Czorlich et al. 2018, Erkinaro et al. 2018). Such changes in demography are likely the result of bottom-up changes in prey community structure, likely fuelled by climate-induced changes to the ecosystem (Frederiksen et al. 2006, Todd et al. 2008).In this study, we demonstrated that the inter-individual variation in diet specialization is linked with age structure as well as the genetic variation in *six6* and *vgll3*, two genomic regions with substantial influence on life history variation and population divergence. This heritable intraspecific variation in diet specialization likely plays an important role in salmon life history by both promoting the niche breadth of species and enabling evolutionary responses in populations to changes in food composition. Given that both genomic regions are highly differentiated among populations, evolutionary response and the resulting demographic trajectories likely differ among populations, concordantly. Future work should focus on characterizing the underlying physiological and/or behavioural mechanisms linking genetic variation with salmon diet acquisition to better predict the evolutionary response of populations to changing environments.

## Data Availability Statement

Data will be available to public in Dryad depository upon acceptance.

## Acknowledgments

We acknowledge the Atlantic Sea salmon fishing organisations in Finnmark county, and especially fishers who collected more than 2000 stomachs from wild Atlantic salmon used in this study.

Ursula Lönnqvist is thanked for laboratory assistance. This project received funding from the Academy of Finland (project numbers 318939 to TA, 307593, 302873 to CRP), from the European Union, Kolarctic ENPI CBC project “Trilateral cooperation on our common resource; the Atlantic salmon in the Barents region (KO197)” (https://prosjekt.fylkesmannen.no/Kolarcticsalmon), and The Research Council of Norway project no. 244086/E50. The funders had no role in study design, data collection and analysis, decision to publish, or preparation of the manuscript.

