## Supplementary material for "Life history genomic regions explain differences in Atlantic salmon marine diet specialization": Supp.

Supplementary Figures (7) and Tables (11)
for

### Life history genomic regions explain differences in Atlantic salmon marine diet specialization

**Supp. figure 1:** Distribution of total and average prey weight in the stomach content dataset (n=2121) before and after transfromation using 10g and 5g bins, respectively.

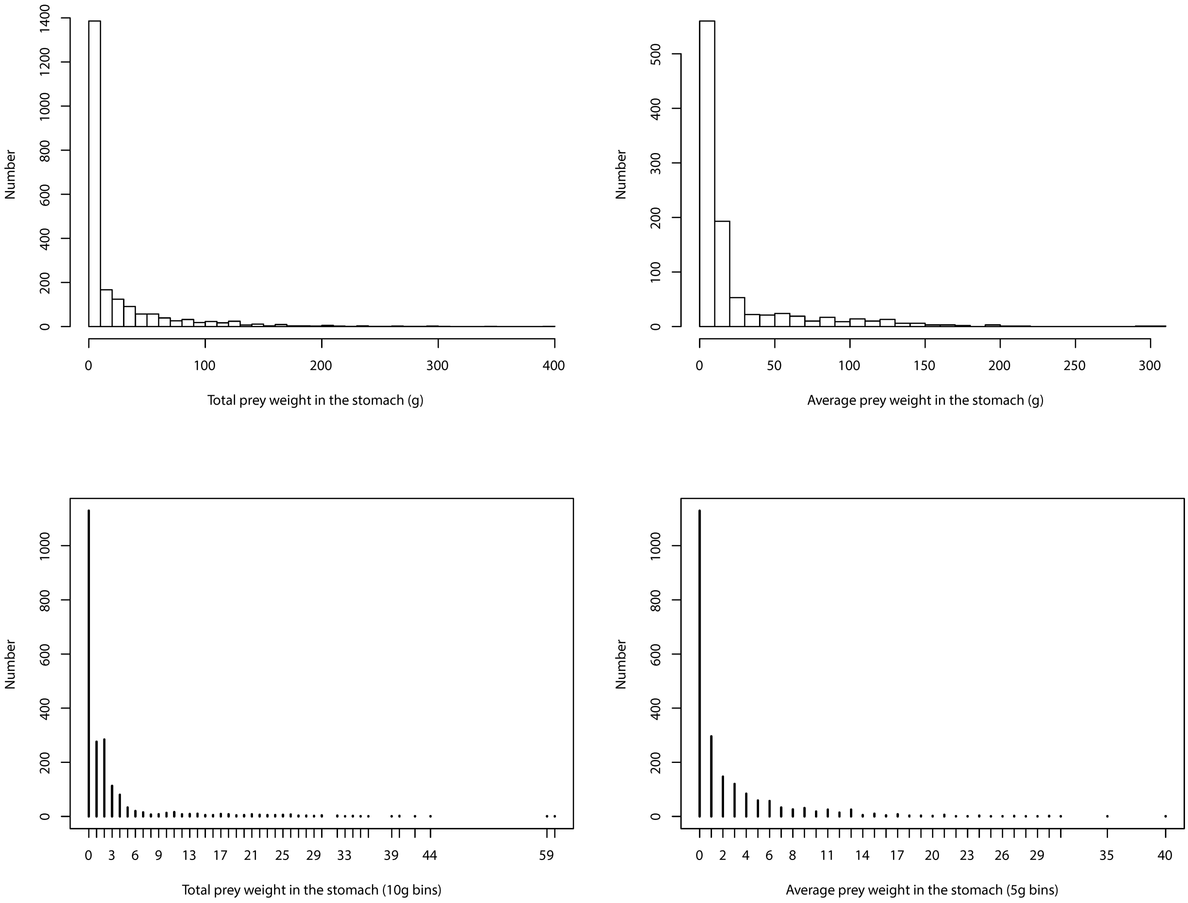

**Supp. figure 2:** Atlantic salmon stomach content in relation to a) prey species number per stomach, b) total weight of prey species , c) Average prey speies weight. All comparsions in (c) are significantly different at p< 0.001 after Tukey comparsion.

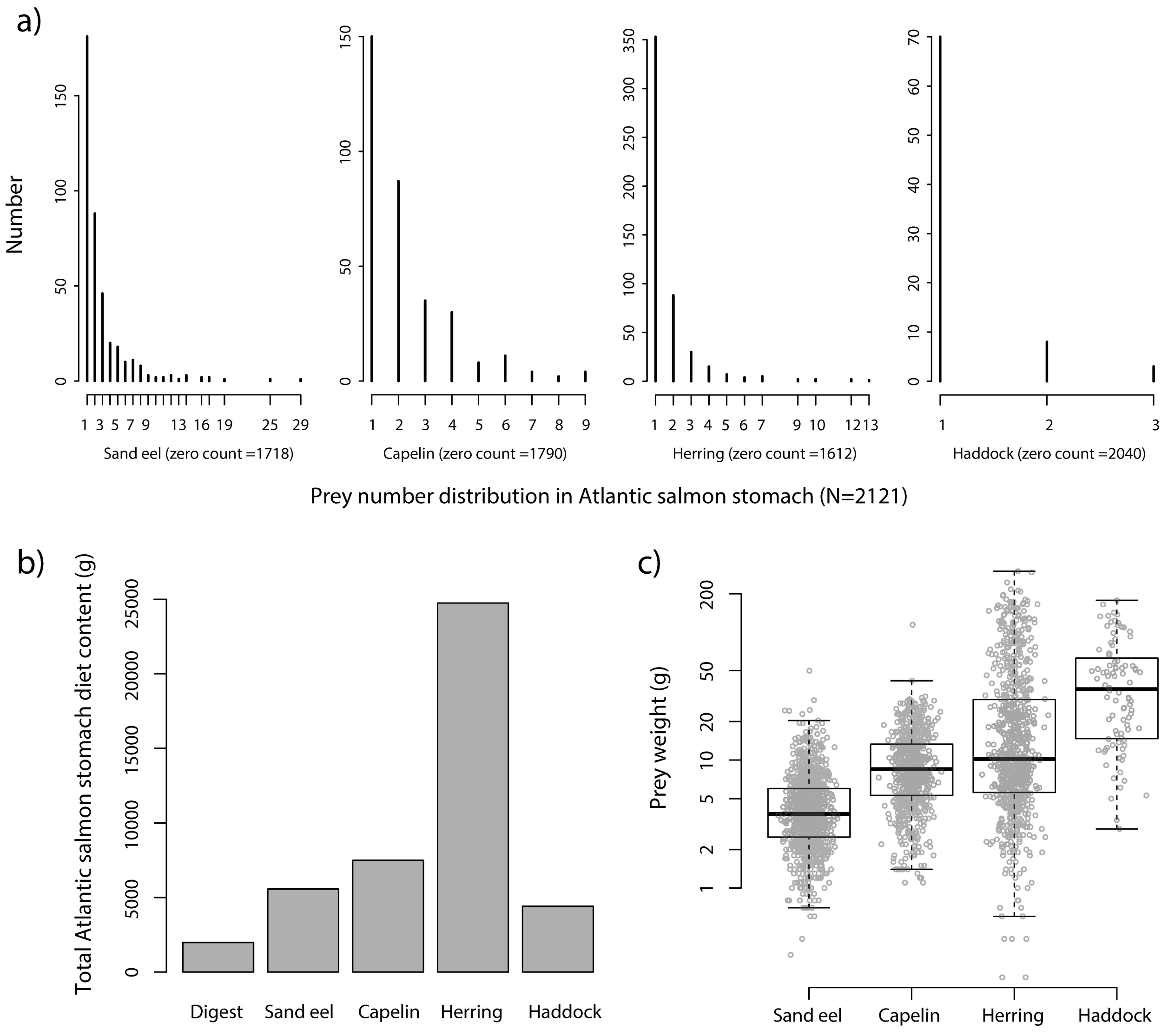

**Supp. figure 3:** Marginal spatio-temporal effects explaining total stomach weight variation in Atlantic salmon, using the two-component hurdle model (e.g. Supp. table 2). Note that the zero-inflation binomial coefficients were reversed for coherent interpretation. In all models, higher cumulative beta coefficient is associated with higher the likelihood of prey presence and quality in the stomach.

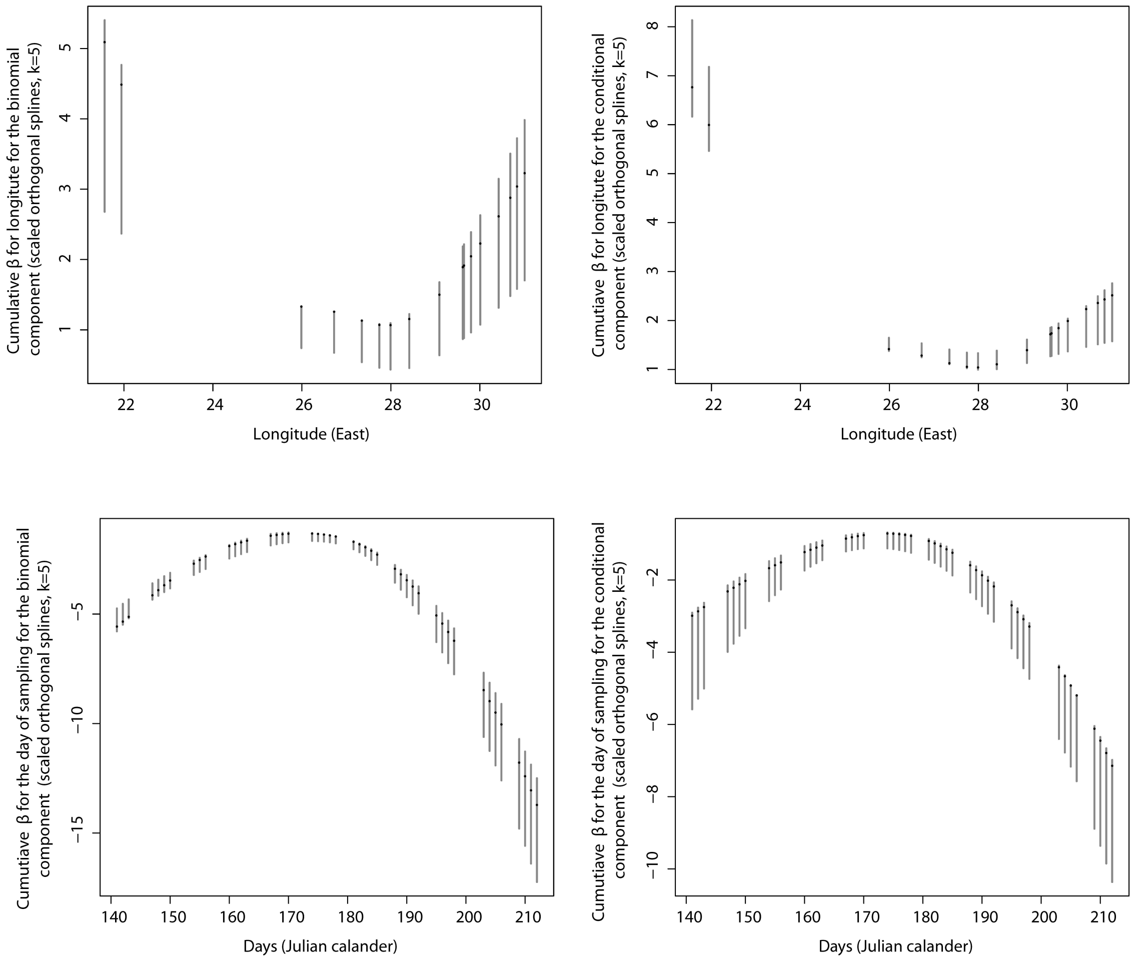

**Supp. figure 4:** Comparing the fit of genetic model (two component hurdle model using total prey weight as the response variable) in the life history genomic regions (*six6* and *vgll3* ) to putatively neutral SNPs in the SNP panel (N=168). The left panel shows the significance of improvement in the genetic model compared to the null when *six6* (vertical red lines) versus other SNPs included in the model. The right panel is same as the left, but for *vgll3*.

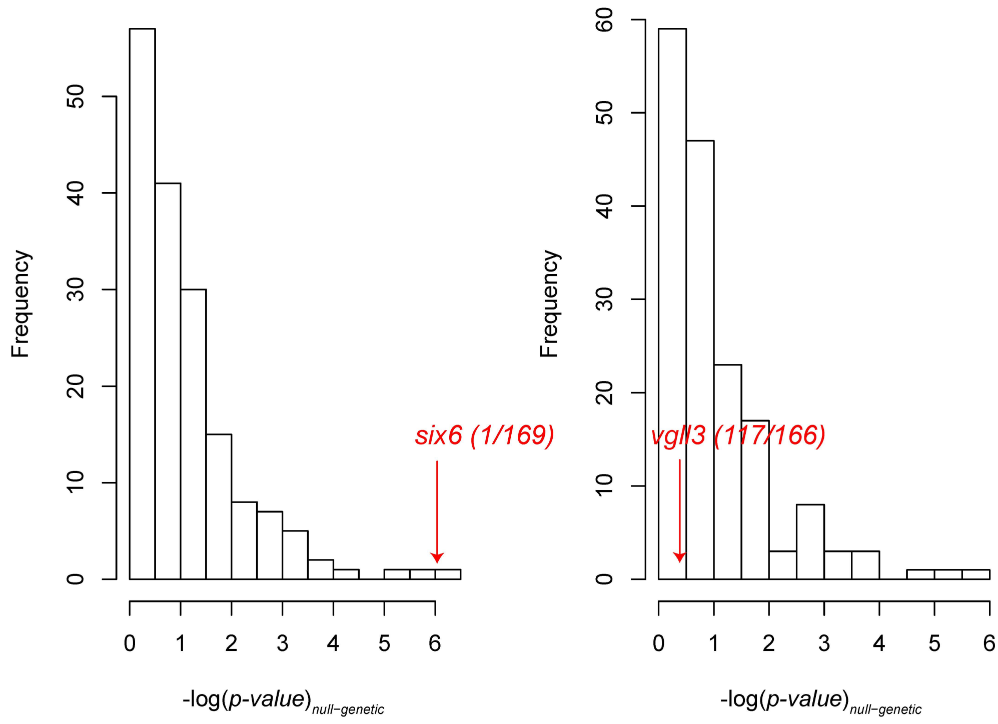

**Supp. figure 5:** Prey preference, measured as the proportional contribution of each prey species to the total stomach content weight, in relation to length at age.

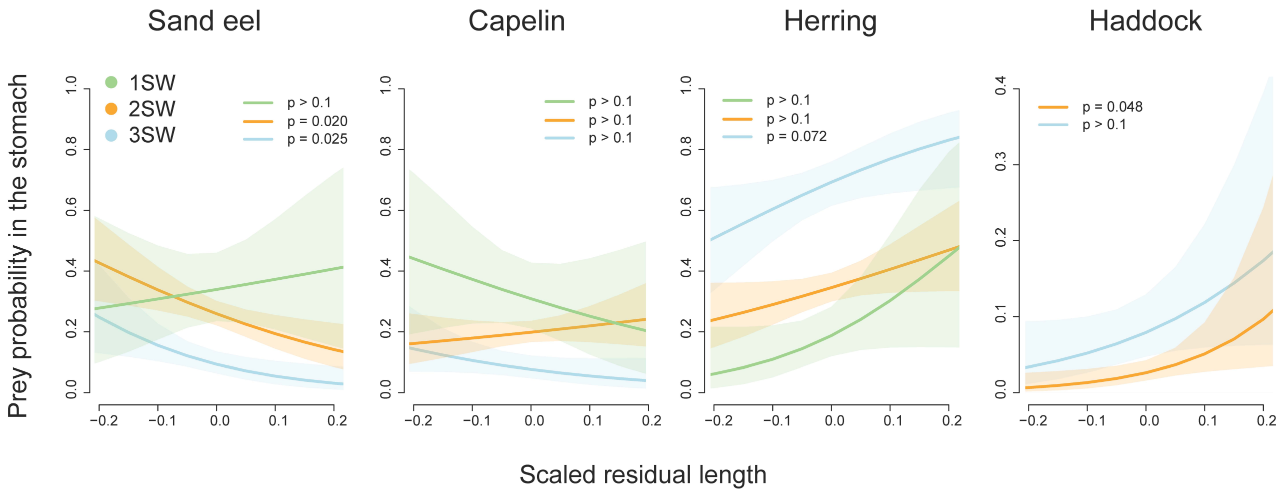

**Supp. figure 6:** Prey preference, measured as the proportional contribution of each prey species to the total stomach content weight, in relation to allelic substitution in life history genomic regions (*six6* and *vgll3*). The dashed line indicates the null situation of no difference in preference between the E and L alleles

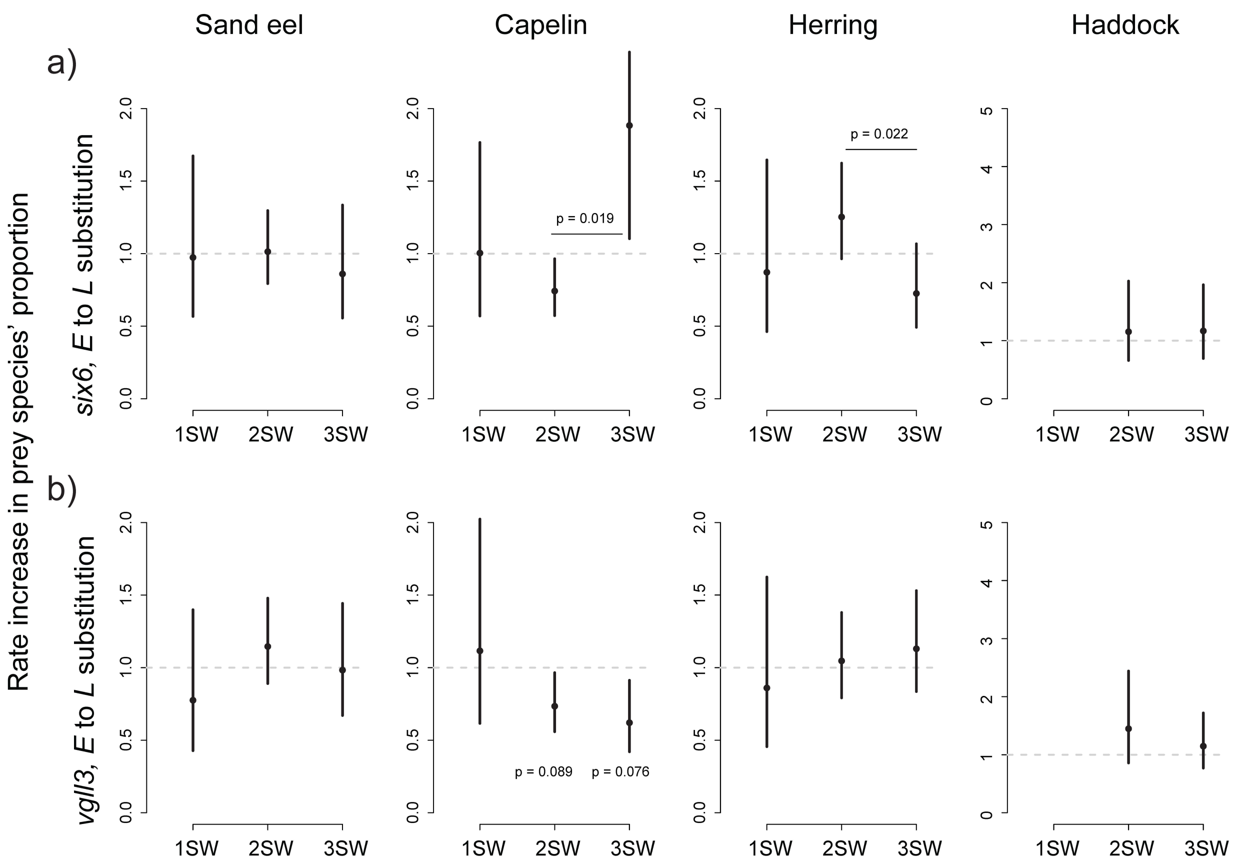

**Supp. figure 7:** Comparison the fit of genetic model (binomial model using species' proportional contribution to total stomach weight as the response variable) in the life history genomic regions (*six6* and *vgll3* ) to putatively neutral SNPs in the SNP panel (N=168). Upper panel shows the significance of improvement in the genetic model compared to the null when *six6* (vertical red lines) versus other SNPs included in the model. Lower panel same as upper, but for *vgll3*.

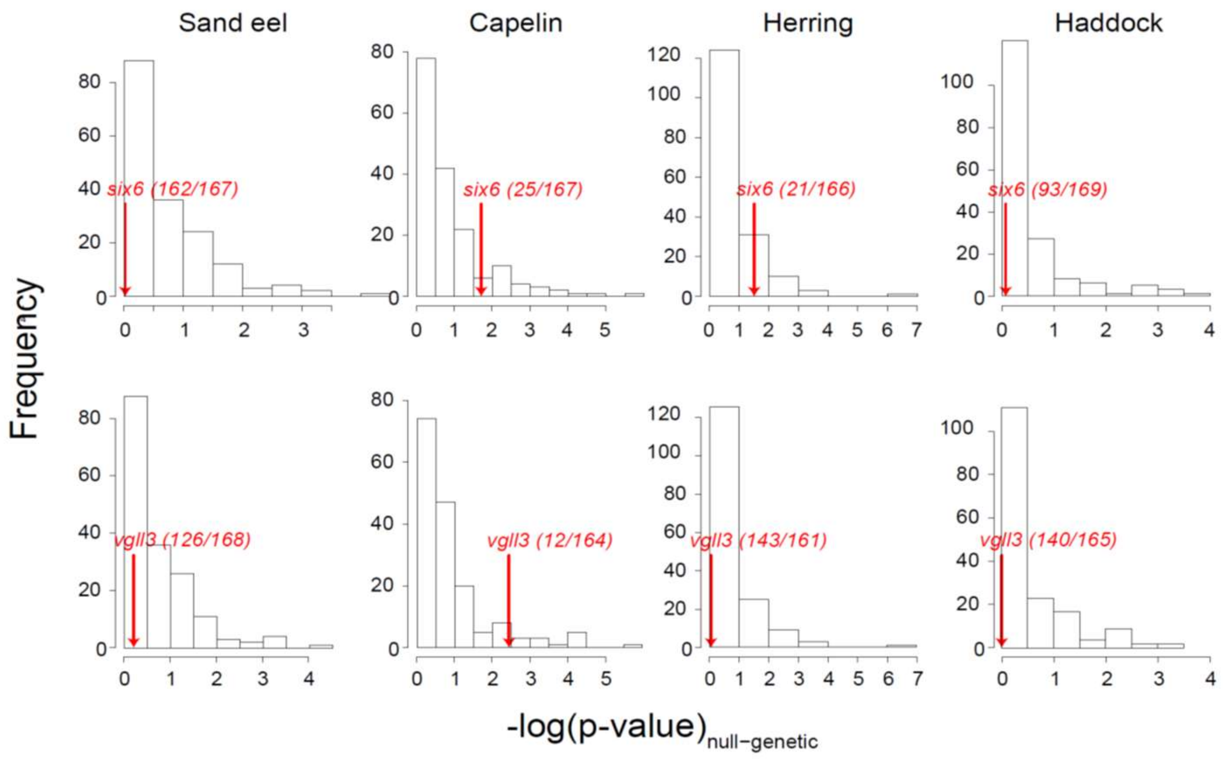

**Supp. table 1:** Primer seqeunces and concentrations for genomic regions amplified with the SNP panel. Refseq refers to Atlantic salmon reference genome assembly ICSASG_v2 (GCF_000233375.1).

| Primer name | Forward primer | reverse primer | refseq chromosome number | refseq start position (bp) | refseq end position (bp) | final primer conc. (microM) | notes |
|---|---|---|---|---|---|---|---|
| akap11_Val214Met_missense | CTTGTCTCTTTCCCTAGCCAAT | GATAAATCCTTGTTTCTCTTGTGG | 25 | 28720727 | 28720825 | 0.05 | |
| vgll3Met54Thr misense | CTCCCTCTCCCTCTCTCTCG | CCCTGGAAACTGCTGCTC | 25 | 28656019 | 28656120 | 0.03 | |
| vgll3Asn323Lys misense | TGTGGGACACAGCACACAC | GGCCTGCTCCACCTCTGT | 25 | 28658085 | 28658169 | 0.03 | |
| six6.top.LD | TCTGTTGCTTGTGTTTGTGTGT | CACAAGTGCCAGGCTAGGAG | 9 | 24904563 | 24904631 | 0.04 | |
| vgll3.top | TCTCCTCTGTTGTCATCCAGAA | ACCCAATCAGACCACACAGC | 25 | 28666870 | 28666943 | 0.08 | |
| SDY (sex) | GCGAAATGAGAGGAGGTGCTTAGTC | GCTTTGGGAGAGAGATGACG | NA | NA | NA | 0.08 | |
| GCR_cBin26140_Ctg1_118 | CCACATCCCTGTCATTGGTA | GCAGATTACACACTAGCCATCG | 1 | 92885957 | 92886051 | 0.04 | |
| ESTNV_30347_1246 | TCATTCTCTGGACTTCACCTCA | AACGGAGAGTCGAACAGAAGTAA | 1 | 148637873 | 148637986 | 0.06 | |
| ESTNV_37143_1231 | TCAAACAATTACAGTCCCTGAAGA | TTCATTTGGCAAGAAACATCTC | 2 | 14031730 | 14031818 | 0.08 | |
| GCR_cBin22155_Ctg1_66 | GCCTACCCTGCTTACTCCTTC | GCCCATAGCTTGACATCACA | 3 | 39855197 | 39855295 | 0.05 | |
| GCR_cBin25322_Ctg1_126 | TGCTACCATCTTTGCCACTTC | TGTGACCCTAACTCGCATTCT | 3 | 63818082 | 63818186 | 0.04 | |
| ESTV_15804_172 | GAGCCCAACATCCCAAGTTA | TGCACAGACCCAGAGTTTCA | 3 | 71053538 | 71053641 | 0.04 | |
| ESTV_15528_560 | GCAGTAGCTTGTCCCATCTCA | TGTAAAGCCTCTGACCGGAAT | 4 | 37204901 | 37205014 | 0.04 | |
| ESTV_17539_1123 | CCCGCCCAGAAGTAAAGAGT | GTTGGCTCAATGGCACATAA | 4 | 61721160 | 61721240 | 0.06 | |
| ESTNV_35811_674 | GCGACCAAATTCAAGAGGAA | ACGATCCCATCAATCTCCAG | 5 | 16014230 | 16014318 | 0.05 | |
| GCR_cBin12244_Ctg1_232 | TTTGTGTAAAATTCGGGTTTCTC | AAAAGGGTTAATGCCCCAAG | 5 | 25340334 | 25340412 | 0.04 | |
| GCR_cBin4585_Ctg1_148 | GCATTGACACACACATTAGCC | GCCCCTTGTCTTTGTTCCTA | 5 | 58425438 | 58425555 | 0.06 | |
| GCR_cBin21411_Ctg1_326 | TGCTGGTGTCCTACATATAACGA | GCTACCCTGAGTCCATTCCA | 5 | 75478092 | 75478196 | 0.04 | |
| ESTV_16910_318 | AGGCAGTCAAAGAGCACCAT | TGCACAAGTATTCAGCAAACAA | 6 | 54237180 | 54237259 | 0.05 | |
| ESTNV_25964_603 | ATCAACCTCCATTAGAAAATGTGAT | AATCTACTTTCAGGCACCGTTT | 10 | 25966905 | 25966989 | 0.06 | |
| ESTNV_34873_1803 | GACGGTGCCCATGATTAAGT | TCTCTTGAGCCGAGTGTGAA | 10 | 66175910 | 66176008 | 0.05 | |
| GCR_cBin8941_Ctg1_434 | TCCCAGTTCAATCAATCAATCA | CGCTGAAGTTGCCTATCAATC | 10 | 91439334 | 91439423 | 0.1 | |
| GCR_cBin3562_Ctg1_54 | TCACTTTCAGCTTGATGTATTATCG | AACACTTGACCTGCCTCCAG | 10 | 112128739 | 112128844 | 0.05 | |
| GCR_cBin35201_Ctg1_151 | GGATGCCTTAGTTTCCACGTA | TGGGAAAGGTTGATTATGTAACTG | 11 | 32973673 | 32973780 | 0.05 | |
| GCR_cBin5691_Ctg1_70 | GGCCACCAATATATACACTAAGTCC | AACCAGTCAACTAATAGTCACTCTGC | 9 | 104664772 | 104664892 | 0.04 | |
| ESTV_17739_134 | CATCATTCCAGGCCACATTT | ACGCAGGACAATCCCATTT | 10 | 39996522 | 39996641 | 0.04 | |
| GCR_cBin23617_Ctg1_247 | CGGCTTCAAATGACTGACC | CACCGCCTGAATCATTTCTAA | 11 | 39312089 | 39312197 | 0.05 | Also amplifies paralog locus |

| | | | | | | | |
|---|---|---|---|---|---|---|---|
| GCR_cBin3754_Ctg1_259 | TATCATGTACAGGTTACCCCATTG | AAGACCTGTCATTTTGTGACCA | 9 | 24123060 | 24123134 | 0.06 | |
| GCR_cBin25404_Ctg1_420 | TGGGTGGGGACATAATACATTT | GGACGTCTCAGAGCGATCA | 16 | 81864081 | 81864191 | 0.05 | |
| GCR_cBin16182_Ctg1_117 | AGTGGGCAGAAATTGGAAACT | TGATCACTGAGGATTTGAGTGC | 20 | 48299071 | 48299175 | 0.04 | |
| GCR_cBin14896_Ctg1_214 | TGGATTTAACCTTTCGAGTTTCA | TGAACTGTATGGCGGTGTCTAT | 16 | 76689689 | 76689767 | 0.08 | |
| ESTV_15429_132 | GACAAATTGCCATTAACATTGC | CTTTGGAAAACATGGGACAGTT | 9 | 25111194 | 25111280 | 0.05 | |
| BASS122_B7_C07_415 | CAGCAAAATCAAATGGTTTACAGT | GATTACATGGCCATTTTCTTTGA | 14 | 66299471 | 66299554 | 0.08 | |
| ESTNV_34763_2153 | AGATGTGTTGCTGGGAGAGATT | ATTGGGTGGATAATTTGAGCAC | 6 | 11402887 | 11402983 | 0.04 | Also amplifies paralog locus |
| ESTNV_28052_855 | CACTCCTATCCATACAGCATTTTTC | TTATACTGTTGGCAGTCAGTTGG | 10 | 21695550 | 21695624 | 0.08 | |
| ESTV_17233_535 | TTCTTAGCGTGGTCAGACTGTT | TCATGTGAAGGACCAGCTAAAC | 12 | 308167 | 308253 | 0.04 | |
| ESTNV_27081_431 | AAGAAGGAGATAGTTGGGCTGA | CTCCAATTGGTTGTTTTTGTTG | 18 | 47612286 | 47612391 | 0.05 | |
| ESTV_16673_1575 | TGTAGCACCATCGTATACTCTTCTGT | CACTGGCTTGATGACTCCTGTA | 1 | 135996047 | 135996165 | 0.04 | |
| GCR_cBin9783_Ctg1_69_V2 | TTCTTGTGTAGGCTGCTTTCCT | GGTCACGTCGCCAGAGTTAG | 1 | 94968741 | 94968825 | 0.04 | |
| GCR_cBin50777_Ctg1_237_V2_NullAll | AGTGCATGACAATGAGCCAGT | AAATAAATCTTTCAGTGGCACACA | 19 | 51819393 | 51819469 | 0.04 | |
| ESTNV_33390_1703 | GCCAACAACCTTCATGTCCTA | GGGAAACCCGTCATTTACAAC | 1 | 56853212 | 56853308 | 0.05 | |
| ESTNV_35387_1829 | TGGACCTTTGGTGTAATACTCG | CATTCAAGTTAAACCCTTCAGCA | 3 | 22742354 | 22742428 | 0.05 | |
| GCR_cBin2723_Ctg1_244 | ATACCCCTTTGGGTCACCTTAC | GGGGTGGTACGGTTGTAACTAT | 3 | 6891775 | 6891856 | 0.05 | |
| GCR_cBin2889_Ctg1_168 | GCCCTGCAGTATTGAGAAGGTA | CCAGTGTCGATTGTTTGATGTT | 29 | 6135214 | 6135300 | 0.06 | |
| GCR_hBin26013_Ctg1_125 | GATACAAGTTCTGCTGCAAAGG | ATCGATCATGTCCTTCCTCAGT | 24 | 6761108 | 6761199 | 0.04 | |
| ESTNV_31116_191 | GAAATAAACATTTTGCCATGGAT | TTTAGATTTCCCGCAACAGACT | 1 | 49611630 | 49611737 | 0.05 | |
| BASS129_B7_H01_304 | ATAGCCAGCACCATATTGCCTA | AGGTCCTTTTAATGCCCATAGC | 9 | 57180431 | 57180505 | 0.03 | |
| ESTNV_19750_154 | CTCCTCTCTCTGGGGAGAAAC | TCTCATACATCCCATCATGCTT | 5 | 34634457 | 34634538 | 0.05 | |
| GCR_cBin21225_Ctg1_149 | TAACTTTCCCCCAAAAGTTCCT | CCACTGAAATTGTTGGAAAAGG | 9 | 110397654 | 110397728 | 0.04 | |
| ESTNV_34745_1760 | TCTGAGATGCTACTCATTGTCCTC | GCCTACAATCAGGTGTCAGTTG | 10 | 76509146 | 76509231 | 0.04 | |
| GCR_cBin10533_Ctg1_181 | CAAGGAAACGTTGGCATTAGAC | CGCAACGACTTCTGTGGTAGTA | 18 | 49453790 | 49453864 | 0.05 | |
| GCR_cBin12749_Ctg1_233 | AGGCACGCATATCTTTAGCAAG | ACGTAAAACATCTGCCTTCTGC | 1 | 64640514 | 64640595 | 0.04 | |
| ESTNV_25560_280 | TAGCCTCCAACAACAAGTCTGA | AAATTAATCGGAGCGACACTTG | 6 | 30067945 | 30068031 | 0.05 | |
| GCR_cBin38774_Ctg1_204 | GGGTTGAGTAGGGCTCACAG | CCACCTTCACTCTGTCGTTG | 1 | 34789554 | 34789669 | 0.03 | |
| ESTNV_36697_1478 | ATCCATCTACCGCTCACTTCTT | CCAACATGCCCTTGGAGA | 2 | 42015090 | 42015183 | 0.1 | |
| BASS124_B7_D10_265 | AAGAAACCCGAGCCCATTT | TGTAAGGGATTGCTGTATTGGA | 3 | 13992860 | 13992945 | 0.04 | |
| GCR_rBin10198_Ctg1_113 | TCAAGGTCAATGCGTCTCAC | CGGTGGATGCCTAAGTCG | 4 | 4114455 | 4114557 | 0.04 | |
| GCR_cBin8054_Ctg1_165 | AGGGAGTATCTTACCCTTTATCTGG | TGGAGAGGGATCAGTCACAAA | 9 | 51922296 | 51922388 | 0.06 | |
| ESTV_15468_87 | CATTTAACCCTTAGCTGGTATGC | GGGAAGAAGCCCTCGAAGT | 12 | 29096295 | 29096413 | 0.04 | |
| GCR_cBin21967_Ctg1_139 | AAACTTACCGGACGTGGAAA | GCAATTCCGAGATGGTTGAC | 12 | 42137069 | 42137186 | 0.04 | |

| | | | | | | | |
|---|---|---|---|---|---|---|---|
| ESTNV_16810_167 | TGTATGTGTGCCATGCTATGTG | TTGTGTCAATGTTGCTACTGTTTC | 12 | 64524836 | 64524919 | 0.05 | |
| ESTNV_36929_2048 | TGAGGAGGAGCAGAGAGAGC | GGAAGTCCCTGGTGGTCTATG | 12 | 90073284 | 90073372 | 0.03 | |
| ESTNV_33495_537 | TGCCAGTAAGGGTCAGAGGT | CAGGGCAGTGTGAGTGGAC | 13 | 2587953 | 2588046 | 0.02 | |
| ESTV_17113_1162 | CCCAGTGAGGTTGGTCAGTAG | TCCAAGGGTGTGAGTGAGTG | 14 | 10538172 | 10538255 | 0.03 | |
| GCR_cBin17206_Ctg1_260 | AACTTGATACGGGCAATGAAA | GCTATATGTCCAATTCTGCCAAT | 14 | 17688158 | 17688259 | 0.05 | |
| ESTV_21094_315 | AGAAGTTGACCGGACGGAAT | CCCTCTCCCTCACCCATC | 14 | 53565940 | 53566029 | 0.04 | |
| ESTNV_33971_297 | GCCGTAATATCGACAAAGCAA | CAACCCGCACCTTTAATTTC | 15 | 64684850 | 64684933 | 0.05 | |
| GCR_cBin27281_Ctg1_182 | GGGAAGTTTGTGGTTGGCTA | TCAGTGTGACTAGAGCCCAAATAA | 16 | 4925233 | 4925348 | 0.04 | |
| GCR_cBin1040_Ctg1_440 | AGCCCAGTCAGGCAGCTA | AATCCGTCAATCATGCCAAC | 17 | 33214851 | 33214926 | 0.03 | |
| GCR_cBin9171_Ctg1_119 | GGCTACCTCCACTTTGGTGA | CCATGTTCCTGTTTATGTGACG | 20 | 69091937 | 69092031 | 0.04 | |
| GCR_cBin17597_Ctg1_113 | GACTCTGGGTGGCTGGATAC | GACCATCACAAGTGCTCTGC | 26 | 16201943 | 16202024 | 0.04 | Also amplifies paralog locus |
| GCR_cBin13983_Ctg1_109 | TTCACAACACAAACAGGGAATTA | TCATCCATTTATCCTTGGCTTT | 26 | 17671512 | 17671631 | 0.08 | |
| GCR_cBin48270_Ctg1_181 | AGAAGGTGAAGCCCAAAGC | TTAAGTAATCTGCCCAACCATGT | 28 | 28235772 | 28235889 | 0.04 | |
| ESTNV_36853_78 | GAACAGGACAGGGAGAGACG | CGTCCGAAGAATATAGGAGTGTG | 1 | 20223405 | 20223493 | 0.03 | |
| GCR_cBin20942_Ctg1_63 | AACAAGGATAGAAATTGGAAGTCTG | GGCAGCAGAGGTCTCACG | 2 | 48412094 | 48412178 | 0.04 | |
| GCR_cBin25027_Ctg1_137 | TCCTCCCATCCTAGAGAGACC | CTGTGTTCTCAGCTACCTCTGTG | 2 | 64471829 | 64471932 | 0.03 | |
| GCR_cBin30698_Ctg1_184 | AAGAGAGTGCCCAGAAATCG | TTGCATGGGTTTAGAGGTGA | 6 | 42334276 | 42334357 | 0.04 | |
| GCR_cBin31213_Ctg1_147 | CTTCTGTTAGGTATTTCTTTGGCTTT | TGCTGTCATGGAATGTAAGGAC | 7 | 47745535 | 47745650 | 0.05 | Also amplifies paralog locus |
| GCR_cBin26895_Ctg1_24 | ATCCTTGCTGCTCCATGC | CATTTAGCGAGTCCTGTATAGTGAAA | 9 | 50963623 | 50963701 | 0.08 | |
| GCR_cBin46328_Ctg1_67 | CCATATTACCAGAGCTTCATCTTATTC | TCACCATCTATTGTTACCTCGAAA | 9 | 107686908 | 107687014 | 0.1 | |
| ESTV_14649_80 | GCCATTCCAGTCAGTCACAA | GGAACAGCTCTCCTTCATTCA | 10 | 17381779 | 17381898 | 0.06 | |
| GCR_cBin12488_Ctg1_395 | TGGCGTATTGTCAGGTCATT | GAACTGGTACGATTAGACAGGTTACA | 10 | 97401480 | 97401566 | 0.1 | |
| ESTNV_25399_259 | GGAGGATAGGATACCGATAGCA | TTGATGTAGGCAGGTCTGTAGG | 17 | 17923767 | 17923849 | 0.04 | |
| ESTNV_29790_168 | ACAACTGGGCACACAGGAG | GTCCAAGGGCTGTCACCA | 20 | 12832004 | 12832103 | 0.02 | |
| GCR_cBin4719_Ctg1_145 | GGCATGTGCTCAACAACAAA | CAGATTGTGCTCCTTTCCTTCT | 21 | 19278752 | 19278855 | 0.06 | |
| GCR_cBin466_Ctg1_234 | CTATTGCGTTGACATGCACA | TACATGCAGCTCTCGCTTTG | 25 | 12402960 | 12403043 | 0.04 | |
| GCR_cBin11286_Ctg1_158 | CGAAACGGCAACAAAGACA | TGCACAAGCTCAGTATCCATTT | 28 | 1673395 | 1673475 | 0.06 | |
| GCR_cBin4881_Ctg1_143 | GCGTGTGTTTGAGCATGAGT | GGACAAGTCAATATGGCTCTTCA | 28 | 30455965 | 30456066 | 0.05 | |
| ESTNV_18264_285 | GGCAGTCTTTCATATTTCTACAAGG | TGCCACAAGGAAGTGAAGG | 29 | 20183822 | 20183926 | 0.05 | |
| ESTNV_34168_1263 | TTTGTTCATGTGCTGTGCAA | GCCCTTGATACTGCCTTGTG | 10 | 12787472 | 12787575 | 0.05 | |
| GCR_cBin28606_Ctg1_136 | TGAAATGTTGTTGTGGTCTTGG | GTGCAGTGACATGGGATCAG | 26 | 13414442 | 13414540 | 0.03 | |
| ESTNV_30970_433 | CAATTAGTGCAGATGGGTAAGGT | TCCGTTCTTTGGCTCATTTC | 13 | 40599181 | 40599276 | 0.04 | |
| ESTNV_12902_251 | CAGGGTGTTGTGTAAAGTTGGA | GTCAAGCACATGGTTAATGTCAG | 4 | 22803310 | 22803392 | 0.06 | |

| | | | | | | | |
|---|---|---|---|---|---|---|---|
| GCR_cBin26759_Ctg1_213 | AGGTCAGCCATCTGAACCA | ACACTTTCACACCACATAAACCA | 10 | 80536845 | 80536962 | 0.04 | |
| ESTV_18792_289 | ATCCCTGTGTTGACCTTTCTG | GATTCTGGCGAGTGGGAAT | 26 | 9248965 | 9249058 | 0.04 | Also amplifies paralog locus |
| AY388592_a | TGTATAAAGTGAGCTGAAGAACTGG | GGGTGTTCCAAAGGCTGA | 11 | 14683805 | 14683880 | 0.06 | |
| GCR_cBin320_Ctg1_314 | TCTTTGGTCTCTGTCCCAGTC | ACATACCTTTCGCGTTCCAC | 19 | 72458660 | 72458744 | 0.04 | |
| GCR_cBin7800_Ctg1_739 | CCATTTCATCAGAGCGATCC | TCAGGTCAGTTCCAGTGTGC | 1 | 157505833 | 157505929 | 0.02 | |
| ESTNV_36166_1693 | AATACTGTTTACCTGTTCCACCAAA | TCCCAGGACAATGGACTAACA | 10 | 40188936 | 40189010 | 0.06 | |
| ESTNV_27052_377 | AAGACCATTTGATTGGCATGT | TCTCTATAATGTTTGTACTGGGACGA | 22 | 32764696 | 32764784 | 0.06 | |
| GCR_cBin28870_Ctg1_219 | ACTCACCTGCCATCCTGTTC | AGATTCCATGCTCAAGTGTAAGG | 6 | 71075560 | 71075666 | 0.06 | |
| ESTV_18550_697 | TTCTACATGCGCCATTCTCA | CGATGACTCCGCCTACCTAGT | 15 | 82875456 | 82875548 | 0.05 | |
| ESTNV_35759_1059 | ATGTGGCAAGAATGCTCCA | GCTCTAGCTCATTTGTGATGATTG | 27 | 21837766 | 21837852 | 0.06 | |
| GCR_cBin7756_Ctg1_139 | CATTCAGACGCAGAACATGG | CTGTCGATGCCTGTCCATTA | 7 | 13950468 | 13950545 | 0.06 | |
| ESTV_13344_491 | CCTCGCTGTGACCCTGTTA | CCTTCCCGTTACCAGTATGC | 1 | 149793899 | 149794018 | 0.04 | |
| ESTNV_29346_496 | AGTGCATGTCCTCTCACCTGT | TCTCTATGGATCGTTGTTTCTCTG | 19 | 51560689 | 51560780 | 0.05 | |
| ESTNV_14089_241 | GTCAACGCATCAGCAGACAC | CAACCTCCTGCACACATCAC | 5 | 30082779 | 30082856 | 0.04 | |
| GCR_cBin1523_Ctg1_267 | TTAGGCCATGCAGTATCCAA | CGACTACTATTGACCGACAGCA | | | | 0.04 | Also amplifies paralog locus |
| GCR_cBin29470_Ctg1_217 | AGAGGCATGTGTTATTGAGTCG | CCAAGTCAACCGCATTGTAA | 22 | 50533547 | 50533638 | 0.05 | |
| GCR_cBin30450_Ctg1_138 | CTTGGGCTTAGGTGAACTCG | CATGAAGGTGTGGTCGTCAG | 14 | 16131455 | 16131558 | 0.03 | |
| ESTV_12100_413 | GCCCAGGTCTGAATGTCTGT | GGCACTGGTTGTGTGTCTTC | 12 | 22289427 | 22289510 | 0.05 | |
| GCR_cBin11531_Ctg1_130 | GTGTTTCTGTTGCTGCTTGG | TTTCAGTGGTGAGGCTGATG | 10 | 75487830 | 75487906 | 0.03 | |
| ESTNV_20594_1113 | CAGTGAGGAGGATAGGATTCAGTT | GGGCAGGGTCTGTAGTTCTCT | 28 | 12809997 | 12810098 | 0.05 | |
| GCR_hBin16337_Ctg1_198 | CCAGACACCTCTTCCTCCAG | GCCTAGACCAGACCCATCCT | 20 | 26931966 | 26932055 | 0.03 | |
| GCR_cBin27882_Ctg1_116 | CCGTAAGGAGCTGCAATAAGAT | CCCGCCATGTTAGCACTC | 5 | 23139781 | 23139881 | 0.06 | |
| ESTNV_36824_935 | CGTCATACCTGTGGCTGATG | GGATCTGTGACTGTGCAAGG | 6 | 31242573 | 31242667 | 0.04 | |
| ESTNV_34414_179 | GGTGTTCAAGGCTCCCATC | ATAGGGCTGACGGCTGTTC | 10 | 115422379 | 115422457 | 0.03 | |
| GCR_cBin4844_Ctg1_156 | TTAACTCATCCCGCTTCGTC | GTGTTATGCGTGTGCCATTC | 9 | 32090400 | 32090485 | 0.04 | |
| GCR_cBin16618_Ctg1_172 | CCCTAACACACTTGCTGCTG | CAAAGATGAATCCTACCACTCAAA | 24 | 22448450 | 22448566 | 0.05 | |
| GCR_cBin15158_Ctg1_411 | CTTTACTGAGGGCGATGAGC | TCCGCTCTTTCTTCCTGTCT | 15 | 42183595 | 42183714 | 0.05 | |
| ESTNV_33151_631 | GAGAATACCACTTATGCCTCCTCT | GAGTAACGCACGCCGACT | 9 | 112242639 | 112242754 | 0.03 | |
| ESTNV_16766_113 | GCGAGGTTGACCACTCTGTAA | GGCACATTCTGGGACAGG | 2 | 16584919 | 16585031 | 0.05 | |
| GCR_cBin48350_Ctg1_134 | TCCTTTGATAACTCACTTCATTGTTT | ACTGAACCCTCTCCTATTACCTCA | 9 | 58513288 | 58513380 | 0.1 | |
| GCR_cBin36482_Ctg1_136 | CTCTAGCCTGCTCCACATCC | GGCTGAACAATTCCTCCTCA | 20 | 37693879 | 37693991 | 0.03 | |
| ESTV_16941_298 | TCTTTCGGTCAACATGGACTT | TCCTACAGGGACAGTTACGACA | 24 | 2549545 | 2549649 | 0.05 | |
| GCR_cBin2711_Ctg2_478 | ACATTGGTTCCACACTGATGTC | CAAGTAGTTAAGGGTTCCTCTTTCA | 12 | 5998193 | 5998311 | 0.08 | |

| | | | | | | | |
|---|---|---|---|---|---|---|---|
| ESTNV_27014_209 | AACACAATTCCCGTGGATCT | AAGGATTTCGGGCACATTTA | 5 | 7599378 | 7599482 | 0.06 | |
| GCR_cBin38737_Ctg1_101 | TGACGAGACCCAAACAGACA | TGTAATTGAGTTGCGCTGATG | 1 | 132476715 | 132476827 | 0.04 | |
| GCR_cBin4742_Ctg1_269 | CGGATTGAGCAGGCTTTC | AGCGTTAGACCGAGAGAAACA | 4 | 39507190 | 39507266 | 0.06 | |
| ESTNV_21177_562 | TCTCAAAGTAGCAGCAGACACC | TTGGACTTCTTCTCGATCAGG | 11 | 21413019 | 21413120 | 0.05 | Also amplifies paralog locus |
| GCR_cBin34_Ctg1_596 | CCAGTTCGGTCTCTGTGTGA | AATCAGTTAATGCGCCCAAC | 1 | 125482760 | 125482875 | 0.05 | |
| BASS114_B6A_B06_358 | TGTGTTGGAATGTGATTGCTT | TTCGTTTATTGATCTGTGTGCTG | 1 | 26711851 | 26711946 | 0.06 | |
| ESTNV_35602_289 | AACCATATTAGGCAGGGTTGC | GATGCAAGGAATGTACGGAAA | 27 | 12316008 | 12316090 | 0.04 | |
| ESTNV_36162_2354 | CCTGAGAAGAACGCACAGGT | CAATTCTGGCCTATACCTCCA | 18 | 18324468 | 18324580 | 0.04 | |
| GCR_cBin17810_Ctg1_125 | CACACACATGCTCCTTGTAAATG | ACCGGCTCTTCCTGTAAACTAA | 10 | 24434418 | 24434538 | 0.05 | |
| GCR_hBin32056_Ctg1_111 | GGTAAGCCCACTTTGCAGTC | ACATCCACACCCGAAACATT | 13 | 4917678 | 4917759 | 0.04 | |
| GCR_cBin14765_Ctg1_160 | CAGATCATGCAAACTACCAATCA | TTCATTCCTGGTCAATTCTCG | 29 | 22891404 | 22891490 | 0.08 | |
| ESTV_21801_199 | TGTTGACCACGTTGACTGC | TGCAGAAGCTGATGTTTCCA | 5 | 73146454 | 73146537 | 0.04 | Also amplifies paralog locus |
| GCR_cBin19415_Ctg1_122 | CTTGCCCAGATCACAATCAA | TGAGTGGAAATAAGCAGAGTGAA | 16 | 8390999 | 8391093 | 0.08 | |
| ESTNV_37358_2427 | ACTGGCATGTGTCTCCCTCT | CGCTGGTGCTGATAGAGTTG | 8 | 18279672 | 18279779 | 0.04 | |
| GCR_cBin25975_Ctg1_238 | CCAGGGTGAAATTGGATAAATG | AATTGAGCTGTCTGTTTCTGAGC | 6 | 69237016 | 69237095 | 0.05 | |
| ESTNV_37208_1240 | TATTCCATTGACAGCCACGA | AACATGACGATAGCGATGAGC | 14 | 10964114 | 10964218 | 0.05 | |
| ESTNV_32773_510 | ACTTGAACAGGTTGCCGTTT | GAGTTCGATGGAGGTTACGC | 23 | 43475853 | 43475945 | 0.03 | |
| GCR_cBin9235_Ctg1_89 | CAGGACAGCGATTACTCAACC | TCGGGCGTCTGAAAGAAAT | 12 | 44855125 | 44855208 | 0.06 | Also amplifies paralog locus |
| GCR_cBin4558_Ctg1_213 | TGGCTTCCCATGTATAATTGC | CCTTATTGTTTCCATCCGTCA | 22 | 18971219 | 18971326 | 0.05 | |
| GCR_cBin28234_Ctg1_102 | CGGGCATGGTAGTGTCAAA | TGCTACAGAGAGATAGAAAGACAGAGG | 9 | 125452826 | 125452900 | 0.04 | |
| GCR_cBin12492_Ctg1_183 | GCAAGAGGAAAGAAGAAACATCA | GTCAGAGCCGAGAGTGGTCT | 1 | 94853153 | 94853256 | 0.06 | |
| ESTNV_30302_546 | TAAACGGAAAGCCCAAGAAA | AGCGCGAGGTACTGTGTGTT | 27 | 29678765 | 29678867 | 0.04 | Also amplifies paralog locus |
| ESTNV_32705_189 | TGGGAAATAAGTAAACAAGTGTGG | CCAATGAAGTGATATGGACATTCT | 13 | 75148068 | 75148177 | 0.06 | |
| GCR_cBin44478_Ctg1_102 | CTGTGTCGTAGCAAGATGTGG | ACCTTGGCCTTTCTCAACTG | 17 | 42884819 | 42884923 | 0.04 | |
| GCR_cBin12816_Ctg1_240 | AGCATAGGATAGGTGAAAGCAAA | CCGTCCCTACACATTGATGA | 26 | 7372895 | 7372981 | 0.08 | |
| GCR_cBin5629_Ctg1_134 | TGATCCTGTGCAAATAAGAATGA | TGTGAATGGATATGAGGGACAC | 4 | 35872231 | 35872325 | 0.06 | |
| GCR_cBin15004_Ctg1_154 | GTCCTGCTGGTGTTGTGTTG | GCATCTTGGGTAGAGGATTTCA | 5 | 17062842 | 17062957 | 0.05 | |
| ESTNV_33138_53 | AATTCAACAGCGAGCGAGTT | CGTCCGGCAAATCAGTAATC | 16 | 44265883 | 44265959 | 0.04 | |
| ESTNV_31016_480 | TCAAATGTTGTAGTTCTTCAGTTCG | AGACTGGTGTGGATAAGGAGAGA | 18 | 58050548 | 58050631 | 0.05 | |
| ESTNV_33506_1739 | AAAGGTTTGTCCCAGCCATA | ACAAGTGGCAGACGGGTTT | 13 | 17532945 | 17533025 | 0.02 | |
| ESTNV_28270_603 | ATTTGTTCCAGGCTTTGGTG | GCTGCTGTCTGTCTGTCCAA | 1 | 4436853 | 4436942 | 0.02 | |
| ESTNV_32341_483 | ATTAACATTCTAACCTTTGTCATTCG | GGTCCTGAGTTGATCTGTAGTGTAGT | 11 | 14781855 | 14781972 | 0.06 | |
| ESTV_18537_221 | TCCAGCAGGTCATAGATCAGAG | GCTCTTCGAGGTCGGATG | 8 | 539013 | 539087 | 0.04 | |

| | | | | | | | |
|---|---|---|---|---|---|---|---|
| ESTNV_36246_1265 | GCCTTGATTTGATTAGCTCTGG | CAATGATGGACAGTTGTGGTG | 18 | 65998964 | 65999060 | 0.04 | |
| ESTNV_26999_214 | GCTTCAGGATGTATCTGTGATGA | AGAGGGATTGGGATAAAGCTG | 22 | 63183795 | 63183891 | 0.06 | |
| ESTV_18493_1052 | TCTTGCCTATATCTCTGGGACTTT | TCTGGCACACATGACGATAAAT | 6 | 7623016 | 7623135 | 0.05 | Also amplifies paralog locus |
| GCR_cBin30624_Ctg1_252 | GCACTTCTATGCCTGCTAGGTT | TCTGTCTCTATCCCTCTCTTCCTC | 27 | 42079336 | 42079423 | 0.04 | |
| ESTNV_30291_631 | GACCCTGAGACCTGGATGG | AGCAGTGGCTTTCTGGTCAT | 11 | 75878560 | 75878643 | 0.04 | |
| GCR_cBin49589_Ctg1_81 | GGGATTGGGAGAAACTGCTA | GGGATCTTTCCTGGGTCCT | 8 | 11567574 | 11567687 | 0.05 | |
| ESTNV_22611_642 | TTGCACCTGGACACTGATCT | GGGTAGAAGTTGTCAGCCACA | 22 | 26004607 | 26004718 | 0.04 | |
| ESTNV_32815_1072 | AGACTTGTTGTCCTTTGACGTG | TGGAGGAGGTATTTATTGAAGTGA | 9 | 83828692 | 83828802 | 0.06 | |
| GCR_cBin24432_Ctg1_186 | TCCTCCTCGGCACTCTACTC | CGCACAGGTTCACAAAGGT | 17 | 38644170 | 38644284 | 0.06 | |
| ESTNV_23454_358 | AACATATACAGGAAGTGACGATTAACA | ACTGGCTGGAGAACCCTCA | 14 | 89639942 | 89640050 | 0.05 | |
| ESTV_10747_366 | TTGACTTTGTGACCCTGCTG | CTGGTTCCTGTGCTGAAGGT | 3 | 36178664 | 36178778 | 0.03 | |
| ESTNV_17157_615 | CTGCTGACCCTCTGACCTTC | GAATCTTGGCACAACACGTC | 25 | 21668960 | 21669047 | 0.04 | |
| GCR_cBin18412_Ctg1_155 | CCAACAACCAACAACCAACA | TGCGGACAATAGGGTGAGAT | 29 | 38586280 | 38586373 | 0.05 | |
| ESTNV_32112_526 | TTTCAGGTTGTCTTGCATCG | TCTCTGCGAATGTGCCATC | | | | 0.05 | locus not aligned in the refseq. |
| GCR_cBin13678_Ctg1_110 | TGGAACCAATTTCCTGTTTGA | AGCAGCCACGGTTTGAATA | 11 | 91013502 | 91013613 | 0.1 | |

**Supp. table 2:** Coefficients of two-component hurdle model explaining total stomach weight variation in Atlantic salmon. Scaled variables are centered to zero mean, and scaled to one standard deviation.

| | | **Estimate** | **Std. Error** | **z value** | **Pr(>\|z\|)** |
|---|---|---|---|---|---|
| Zero inflation (binomial) model | SW1 | **-0.445624** | **0.1677084** | **-2.657135** | **0.0078808** |
| | SW2 | **-0.074029** | **0.0727647** | **-1.0173752** | **0.308975** |
| | SW3 | **0.4241013** | **0.0982162** | **4.3180363** | **1.57E-05** |
| | scaled (days, knot1) | **0.6850584** | **0.1571405** | **4.3595279** | **1.30E-05** |
| | scaled (days, knot2) | **0.7750237** | **0.1444733** | **5.3644772** | **8.12E-08** |
| | scaled (days, knot3) | **0.7436651** | **0.1604701** | **4.6342896** | **3.58E-06** |
| | scaled (days, knot4) | **0.2046792** | **0.0793964** | **2.5779406** | **0.0099391** |
| | scaled (days, knot5) | **1.3839242** | **0.2321758** | **5.9606734** | **2.51E-09** |
| | scaled (longitude, knot1) | **-0.7940023** | **0.368499** | **-2.1546931** | **0.0311859** |
| | scaled (longitude, knot2) | **-0.4221507** | **0.110282** | **-3.8279213** | **0.0001292** |
| | scaled (longitude, knot3) | **-1.133569** | **0.3457654** | **-3.2784342** | **0.0010438** |
| | scaled (longitude, knot4) | **0.415695** | **0.1048012** | **3.9665116** | **7.29E-05** |
| | scaled (longitude, knot5) | **-0.3063694** | **0.1022574** | **-2.9960623** | **0.0027349** |
| | SW1 : scaled (residual Length) | **0.2120318** | **0.1821991** | **1.1637369** | **0.2445307** |
| | SW2 : scaled (residual Length) | **0.149775** | **0.0677279** | **2.2114233** | **0.0270065** |
| | SW3 : scaled (residual Length) | **-0.0223904** | **0.080516** | **-0.2780862** | **0.7809462** |
| | SW1 : scaled (six6) | **-0.3345** | **0.1760365** | **-1.9001743** | **0.0574103** |
| | SW2 : scaled (six6) | **-0.1661034** | **0.0676199** | **-2.4564284** | **0.0140326** |
| | SW3 : scaled (six6) | **0.0280844** | **0.099784** | **0.2814522** | **0.7783636** |
| | SW1 : scaled (vgll3) | **0.1821643** | **0.2016289** | **0.9034629** | **0.3662803** |
| | SW2 : scaled (vgll3) | **-0.0374022** | **0.0705516** | **-0.5301392** | **0.5960154** |
| | SW3 : scaled (vgll3) | **0.0384426** | **0.075156** | **0.5115039** | **0.6089983** |
| Conditional, zero truncated model | | **Estimate** | **Std. Error** | **z value** | **Pr(>\|z\|)** |
| | SW1 | **0.1739482** | **0.1718191** | **1.0123918** | **0.3113507** |
| | SW2 | **0.9501714** | **0.0857544** | **11.080143** | **1.57E-28** |
| | SW3 | **1.4014971** | **0.105326** | **13.306282** | **2.13E-40** |
| | scaled (sDays1) | **-0.4027841** | **0.16017** | **-2.5147294** | **0.0119124** |

| | | | | | |
|---|---|---|---|---|---|
| | scaled (sDays2) | **-0.349337** | **0.1450009** | **-2.4092051** | **0.0159873** |
| | scaled (sDays3) | **-0.3007373** | **0.1594573** | **-1.8860059** | **0.0592942** |
| | scaled (sDays4) | **-0.1819054** | **0.0752354** | **-2.4178173** | **0.0156139** |
| | scaled (sDays5) | **-0.6693442** | **0.2388199** | **-2.802716** | **0.0050674** |
| | scaled (sLong1) | **1.7134144** | **0.3650874** | **4.6931623** | **2.69E-06** |
| | scaled (sLong2) | **0.5356874** | **0.1190055** | **4.5013665** | **6.75E-06** |
| | scaled (sLong3) | **1.5749021** | **0.333239** | **4.7260431** | **2.29E-06** |
| | scaled (sLong4) | **0.03703** | **0.1005756** | **0.3681808** | **0.7127384** |
| | scaled (sLong5) | **0.1182692** | **0.1118524** | **1.0573692** | **0.2903431** |
| | SW1 : scaled (residual Length) | **-0.2560991** | **0.1724885** | **-1.4847318** | **0.1376149** |
| | SW2 : scaled (residual Length) | **0.1174742** | **0.0562557** | **2.0882191** | **0.0367781** |
| | SW3 : scaled (residual Length) | **0.0405758** | **0.0752636** | **0.539115** | **0.5898075** |
| | SW1 : scaled (six6) | **0.4703079** | **0.1709538** | **2.7510819** | **0.0059399** |
| | SW2 : scaled (six6) | **0.0997124** | **0.0606045** | **1.6452961** | **0.0999088** |
| | SW3 : scaled (six6) | **-0.0132545** | **0.0974867** | **-0.1359623** | **0.8918511** |
| | SW1 : scaled (vgll3) | **0.1206484** | **0.1587451** | **0.7600133** | **0.4472466** |
| | SW2 : scaled (vgll3) | **0.0083112** | **0.0652942** | **0.1272892** | **0.8987115** |
| | SW3 : scaled (vgll3) | **0.0881542** | **0.0738738** | **1.1933084** | **0.2327486** |
| | Random variance | | | | |
| | population | | | | |
| Zero inflation (binomial) model | **6.58E-03** | | | | |
| Conditional, zero truncated model | **0.001741** | | | | |

**Supp. table 3:** Coefficients of the model explaining log(Length) variation in Atlantic salmon. Scaled variables are centered to zero mean, and scaled one standard deviation. Modelling was performed using lmer function in *lme4*, and p values were assessed using the *lmerTest* package in R. Random population term was not a significant source variation.

| | Estimate | Std. Error | df | t value | Pr(>\|t\|) |
|---|---|---|---|---|---|
| SW1 | **4.038817223** | **0.005810102** | **945.7216914** | **695.137108** | **0** |
| SW2 | **4.29347314** | **0.003131384** | **140.0407606** | **1371.110591** | **6.16E-291** |
| SW3 | **4.503492222** | **0.003876493** | **224.7312224** | **1161.743928** | **0** |
| scaled (days, knot1) | **-0.002607727** | **0.005036548** | **2008.597034** | **-0.517760834** | **0.604682166** |
| scaled (days, knot2) | **-0.001790191** | **0.004616246** | **2008.786682** | **-0.387802396** | **0.69820336** |
| scaled (days, knot3) | **-0.003787432** | **0.005116713** | **1995.796265** | **-0.74020804** | **0.459260835** |
| scaled (days, knot4) | **0.0025003** | **0.002553424** | **2034.492876** | **0.979194918** | **0.3276001** |
| scaled (days, knot5) | **-0.000243527** | **0.007298584** | **1994.720482** | **-0.033366278** | **0.973385839** |
| scaled (longitude, knot1) | **0.039982172** | **0.012789822** | **1686.213665** | **3.126093031** | **0.001801675** |
| scaled (longitude, knot2) | **0.008980995** | **0.003786417** | **1606.144453** | **2.371898196** | **0.017814257** |
| scaled (longitude, knot3) | **0.046292597** | **0.011861033** | **1742.231492** | **3.902914542** | **9.87E-05** |
| scaled (longitude, knot4) | **-0.009850401** | **0.003405814** | **1893.937243** | **-2.892231186** | **0.003868821** |
| scaled (longitude, knot5) | **0.003122904** | **0.003800153** | **1052.477125** | **0.821783906** | **0.411386014** |
| SW1 : scaled (six6) | **0.015540114** | **0.005310726** | **2035.019328** | **2.926175041** | **0.003469605** |
| SW2 : scaled (six6) | **0.021420911** | **0.002109138** | **2029.241883** | **10.15623834** | **1.13E-23** |
| SW3 : scaled (six6) | **0.017941362** | **0.003146149** | **2026.954932** | **5.702643039** | **1.35E-08** |
| SW1 : scaled (vgll3) | **0.012824752** | **0.006236594** | **1990.582756** | **2.056371009** | **0.039876871** |
| SW2 : scaled (vgll3) | **0.013351406** | **0.002299867** | **2032.001756** | **5.805294662** | **7.44E-09** |
| SW3 : scaled (vgll3) | **0.014827511** | **0.002370109** | **2007.151506** | **6.256047015** | **4.81E-10** |

**Supp. table 4:** Coefficients of two-component hurdle model explaining total stomach weight variation in Atlantic salmon, when covarying phenotypes, sea age and length, were excluded from the model. Scaled variables are centered to zero mean, and scaled to one standard deviation.

| Zero inflation (binomial) model | | Estimate | Std. Error | z value | Pr(>\|z\|) |
|---|---|---|---|---|---|
| | | 0.1334144 | 0.044798626 | 2.9780911 | 0.0029005 |
| | scaled (six6) | -0.0136844 | 0.047086099 | -0.2906259 | 0.7713375 |
| | scaled (vgll3) | 0.0896929 | 0.046164479 | 1.942898 | 0.0520285 |
| | scaled (days, knot1) | 0.6794616 | 0.152863345 | 4.4448958 | 8.79E-06 |
| | scaled (days, knot2) | 0.7805214 | 0.140569792 | 5.5525541 | 2.82E-08 |
| | scaled (days, knot3) | 0.681098 | 0.155585339 | 4.3776492 | 1.20E-05 |
| | scaled (days, knot4) | 0.1635205 | 0.075877889 | 2.1550484 | 0.031158 |
| | scaled (days, knot5) | 1.2668469 | 0.223774002 | 5.6612781 | 1.50E-08 |
| | scaled (longitude, knot1) | -0.9604613 | 0.35336429 | -2.7180485 | 0.0065668 |
| | scaled (longitude, knot2) | -0.5100245 | 0.104479588 | -4.8815704 | 1.05E-06 |
| | scaled (longitude, knot3) | -1.2596597 | 0.331586398 | -3.7988883 | 0.0001453 |
| | scaled (longitude, knot4) | 0.3455878 | 0.095450015 | 3.6206153 | 0.0002939 |
| | scaled (longitude, knot5) | -0.3455967 | 0.096788308 | -3.5706454 | 0.0003561 |
| Conditional, zero truncated model | | Estimate | Std. Error | z value | Pr(>\|z\|) |
| | intercept | 0.9757788 | 0.084767781 | 11.511199 | 1.16E-30 |
| | scaled (six6) | 0.1493688 | 0.04870415 | 3.0668594 | 0.0021632 |
| | scaled (vgll3) | 0.1068783 | 0.045753014 | 2.3359838 | 0.0194921 |
| | scaled (sDays1) | -0.2598862 | 0.163726191 | -1.5873223 | 0.1124397 |
| | scaled (sDays2) | -0.2190874 | 0.148670553 | -1.4736437 | 0.1405775 |
| | scaled (sDays3) | -0.2101766 | 0.16301288 | -1.2893254 | 0.197285 |
| | scaled (sDays4) | -0.2037484 | 0.07808637 | -2.6092702 | 0.0090736 |
| | scaled (sDays5) | -0.5530733 | 0.24557305 | -2.2521742 | 0.0243113 |
| | scaled (sLong1) | 1.5092333 | 0.368928637 | 4.0908543 | 4.30E-05 |
| | scaled (sLong2) | 0.4466135 | 0.119934182 | 3.7238217 | 0.0001962 |
| | scaled (sLong3) | 1.4451424 | 0.338608702 | 4.2678833 | 1.97E-05 |
| | scaled (sLong4) | -0.067418 | 0.101976219 | -0.6611152 | 0.5085384 |
| | scaled (sLong5) | 0.0318391 | 0.113338274 | 0.280921 | 0.778771 |

**Supp. table 5:** Coefficients of two-component hurdle model explaining total stomach weight variation in Atlantic salmon, after excluding low assignment confidence or imputed individuals (N=1372). Scaled variables are centered to zero mean, and scaled to one standard deviation.

| | | Estimate | Std. Error | z value | Pr(>\|z\|) |
|---|---|---|---|---|---|
| Zero inflation (binomial) model | SW1 | **-0.5805013** | **0.2143019** | **-2.7088009** | **0.0067527** |
| | SW2 | **0.0648402** | **0.0977706** | **0.6631865** | **0.5072111** |
| | SW3 | **0.3983641** | **0.1211693** | **3.2876666** | **0.0010102** |
| | scaled (days, knot1) | **0.7583983** | **0.1898664** | **3.9943783** | **6.49E-05** |
| | scaled (days, knot2) | **0.7459493** | **0.1864663** | **4.0004521** | **6.32E-05** |
| | scaled (days, knot3) | **0.8289446** | **0.20182** | **4.1073473** | **4.00E-05** |
| | scaled (days, knot4) | **0.2795714** | **0.104813** | **2.6673355** | **0.0076455** |
| | scaled (days, knot5) | **1.5737504** | **0.3055734** | **5.1501545** | **2.60E-07** |
| | scaled (longitude, knot1) | **-0.9905804** | **0.472728** | **-2.0954552** | **0.0361305** |
| | scaled (longitude, knot2) | **-0.4210524** | **0.1368899** | **-3.0758469** | **0.0020991** |
| | scaled (longitude, knot3) | **-1.3817125** | **0.4398888** | **-3.1410494** | **0.0016834** |
| | scaled (longitude, knot4) | **0.4545899** | **0.1325038** | **3.4307678** | **0.0006019** |
| | scaled (longitude, knot5) | **-0.2724193** | **0.1295439** | **-2.1029104** | **0.0354736** |
| | SW1 : scaled (residual Length) | **0.0938531** | **0.2441811** | **0.3843586** | **0.7007127** |
| | SW2 : scaled (residual Length) | **0.1419447** | **0.0872878** | **1.6261676** | **0.103914** |
| | SW3 : scaled (residual Length) | **-0.0276343** | **0.0984752** | **-0.2806222** | **0.7790002** |
| | SW1 : scaled (six6) | **-0.323644** | **0.2240759** | **-1.44435** | **0.1486405** |
| | SW2 : scaled (six6) | **-0.206312** | **0.0877126** | **-2.3521351** | **0.018666** |
| | SW3 : scaled (six6) | **0.0534814** | **0.1192164** | **0.448608** | **0.6537144** |
| | SW1 : scaled (vgll3) | **0.4367397** | **0.2755058** | **1.5852289** | **0.1129143** |
| | SW2 : scaled (vgll3) | **0.0029885** | **0.0918863** | **0.0325242** | **0.974054** |
| | SW3 : scaled (vgll3) | **0.0421161** | **0.0908001** | **0.463833** | **0.6427674** |
| Conditional, zero truncated model | | **Estimate** | **Std. Error** | **z value** | **Pr(>\|z\|)** |
| | SW1 | **0.2901495** | **0.2060253** | **1.4083201** | **0.1590363** |
| | SW2 | **0.8844746** | **0.1104487** | **8.008011** | **1.17E-15** |
| | SW3 | **1.4133024** | **0.1155495** | **12.23114** | **2.12E-34** |
| | scaled (sDays1) | **-0.4377122** | **0.187388** | **-2.3358604** | **0.0194985** |

| | | | | | |
|---|---|---|---|---|---|
| | scaled (sDays2) | **-0.4413396** | **0.1826665** | **-2.416095** | **0.015688** |
| | scaled (sDays3) | **-0.3829341** | **0.1934757** | **-1.9792358** | **0.0477895** |
| | scaled (sDays4) | **-0.1624126** | **0.0992368** | **-1.6366168** | **0.1017106** |
| | scaled (sDays5) | **-0.8063697** | **0.3075475** | **-2.6219357** | **0.0087432** |
| | scaled (sLong1) | **1.7528888** | **0.4704769** | **3.7257701** | **0.0001947** |
| | scaled (sLong2) | **0.6492752** | **0.1487906** | **4.3636845** | **1.28E-05** |
| | scaled (sLong3) | **1.6352781** | **0.4229188** | **3.8666477** | **0.0001103** |
| | scaled (sLong4) | **-0.0489063** | **0.1281841** | **-0.3815319** | **0.7028086** |
| | scaled (sLong5) | **0.1871986** | **0.1400734** | **1.3364322** | **0.1814081** |
| | SW1 : scaled (residual Length) | **-0.4820143** | **0.2053734** | **-2.3470146** | **0.0189245** |
| | SW2 : scaled (residual Length) | **0.2016827** | **0.076832** | **2.6249842** | **0.0086653** |
| | SW3 : scaled (residual Length) | **0.0071499** | **0.0874806** | **0.0817312** | **0.9348604** |
| | SW1 : scaled (six6) | **0.2632714** | **0.2067293** | **1.2735084** | **0.2028377** |
| | SW2 : scaled (six6) | **0.1576715** | **0.080373** | **1.9617467** | **0.049792** |
| | SW3 : scaled (six6) | **0.0539914** | **0.1093174** | **0.4938957** | **0.6213799** |
| | SW1 : scaled (vgll3) | **0.2920971** | **0.2044999** | **1.4283484** | **0.1531916** |
| | SW2 : scaled (vgll3) | **-0.0636992** | **0.0866829** | **-0.7348528** | **0.4624292** |
| | SW3 : scaled (vgll3) | **0.0908426** | **0.0906104** | **1.0025624** | **0.3160721** |
| | Random variance | | | | |
| | population | | | | |
| Zero inflation (binomial) model | **8.11E-02** | | | | |
| Conditional, zero truncated model | **0.0417** | | | | |

**Supp. table 6:** Coefficients of two-component hurdle model explaining total stomach weight variation in Atlantic salmon when digested, unidentified material in the stomach content was included in the analysis. Scaled variables are centered to zero mean, and scaled to one standard deviation.

| | | Estimate | Std. Error | z value | Pr(>\|z\|) |
|---|---|---|---|---|---|
| Zero inflation (binomial) model | SW1 | **-1.28654561** | **0.190228576** | **-6.763156384** | **1.35E-11** |
| | SW2 | **-0.6147269** | **0.075289456** | **-8.164847188** | **3.22E-16** |
| | SW3 | **-0.050640692** | **0.089258744** | **-0.567347129** | **0.570478363** |
| | scaled (days, knot1) | **0.711455298** | **0.160083598** | **4.44427354** | **8.82E-06** |
| | scaled (days, knot2) | **0.713266642** | **0.147148901** | **4.847244111** | **1.25E-06** |
| | scaled (days, knot3) | **0.706499693** | **0.164450271** | **4.296129693** | **1.74E-05** |
| | scaled (days, knot4) | **0.151484573** | **0.078971473** | **1.918218903** | **0.05508326** |
| | scaled (days, knot5) | **1.288890254** | **0.233558017** | **5.518501446** | **3.42E-08** |
| | scaled (longitude, knot1) | **-0.481949221** | **0.37729756** | **-1.277371688** | **0.201471055** |
| | scaled (longitude, knot2) | **-0.440088364** | **0.110059163** | **-3.998652665** | **6.37E-05** |
| | scaled (longitude, knot3) | **-0.722867624** | **0.352911484** | **-2.048297254** | **0.040530882** |
| | scaled (longitude, knot4) | **0.297811097** | **0.103202736** | **2.885689947** | **0.003905567** |
| | scaled (longitude, knot5) | **-0.46191972** | **0.101514026** | **-4.55030442** | **5.36E-06** |
| | SW1 : scaled (residual Length) | **0.151684936** | **0.209187131** | **0.725116003** | **0.468380868** |
| | SW2 : scaled (residual Length) | **0.294813937** | **0.07115482** | **4.143274313** | **3.42E-05** |
| | SW3 : scaled (residual Length) | **0.006131676** | **0.078167545** | **0.07844274** | **0.937475876** |
| | SW1 : scaled (six6) | **-0.112221415** | **0.203186924** | **-0.552306284** | **0.580738526** |
| | SW2 : scaled (six6) | **-0.176931092** | **0.070610917** | **-2.505718648** | **0.012220282** |
| | SW3 : scaled (six6) | **-0.029034895** | **0.097150537** | **-0.298864992** | **0.76504306** |
| | SW1 : scaled (vgll3) | **0.022323774** | **0.233273746** | **0.095697758** | **0.923760622** |
| | SW2 : scaled (vgll3) | **-0.035129433** | **0.073158525** | **-0.480182362** | **0.631097727** |
| | SW3 : scaled (vgll3) | **-0.052448859** | **0.073180658** | **-0.716703841** | **0.473556855** |
| Conditional, zero truncated model | | **Estimate** | **Std. Error** | **z value** | **Pr(>\|z\|)** |
| | SW1 | **-0.741469311** | **0.291728804** | **-2.541639022** | **0.011033406** |
| | SW2 | **0.069040359** | **0.237004303** | **0.291304243** | **0.770818642** |
| | SW3 | **0.454493837** | **0.242302519** | **1.8757289** | **0.060692515** |
| | scaled (sDays1) | **-0.484660729** | **0.189578248** | **-2.556520771** | **0.010572476** |

| | | | | | |
|---|---|---|---|---|---|
| | scaled (sDays2) | **-0.487334316** | **0.171872531** | **-2.835440385** | **0.004576255** |
| | scaled (sDays3) | **-0.424765415** | **0.187282899** | **-2.268041652** | **0.023326668** |
| | scaled (sDays4) | **-0.265891839** | **0.08853519** | **-3.00323338** | **0.002671275** |
| | scaled (sDays5) | **-0.960353593** | **0.27855108** | **-3.447674998** | **0.000565434** |
| | scaled (sLong1) | **2.13903141** | **0.43745439** | **4.88972441** | **1.01E-06** |
| | scaled (sLong2) | **0.657306874** | **0.141834821** | **4.634312423** | **3.58E-06** |
| | scaled (sLong3) | **2.096464945** | **0.401719313** | **5.218730784** | **1.80E-07** |
| | scaled (sLong4) | **-0.096787662** | **0.116996197** | **-0.827271858** | **0.408082991** |
| | scaled (sLong5) | **0.135902093** | **0.131354029** | **1.034624472** | **0.300844321** |
| | SW1 : scaled (residual Length) | **-0.413217145** | **0.196269552** | **-2.105355315** | **0.03526039** |
| | SW2 : scaled (residual Length) | **0.139082288** | **0.066041953** | **2.105968752** | **0.035207067** |
| | SW3 : scaled (residual Length) | **0.041456325** | **0.089052952** | **0.465524437** | **0.641555941** |
| | SW1 : scaled (six6) | **0.606417705** | **0.186649806** | **3.248959733** | **0.001158279** |
| | SW2 : scaled (six6) | **0.119392645** | **0.071968577** | **1.658955195** | **0.097124821** |
| | SW3 : scaled (six6) | **-0.031543498** | **0.117508391** | **-0.268436129** | **0.788363633** |
| | SW1 : scaled (vgll3) | **0.129891295** | **0.177242171** | **0.732846445** | **0.46365209** |
| | SW2 : scaled (vgll3) | **0.005724001** | **0.07849863** | **0.072918479** | **0.941870989** |
| | SW3 : scaled (vgll3) | **0.04838387** | **0.087716205** | **0.551595574** | **0.581225469** |
| | Random variance | | | | |
| | population | | | | |
| Zero inflation (binomial) model | **1.52E-08** | | | | |
| Conditional, zero truncated model | **0.03907** | | | | |

**Supp. table 7:** Coefficients of conditional, zero truncated model explaining average prey number in Atlantic salmon diet. Scaled variables are centered to zero mean, and scaled to one standard deviation.

| | | | | | |
|---|---|---|---|---|---|
| Zero inflation (binomial) model | SW1 | **-0.448805298** | **0.167486081** | **-2.679657283** | **0.007369757** |
| | SW2 | **-0.074819871** | **0.072484955** | **-1.032212423** | **0.301972616** |
| | SW3 | **0.413139387** | **0.097702069** | **4.228563336** | **2.35E-05** |
| | scaled (days, knot1) | **0.695137515** | **0.157079029** | **4.425399885** | **9.63E-06** |
| | scaled (days, knot2) | **0.774359136** | **0.144409022** | **5.362262881** | **8.22E-08** |
| | scaled (days, knot3) | **0.751444522** | **0.160421886** | **4.684177093** | **2.81E-06** |
| | scaled (days, knot4) | **0.207608835** | **0.079276279** | **2.618801442** | **0.008823929** |
| | scaled (days, knot5) | **1.389649221** | **0.232016626** | **5.989438107** | **2.11E-09** |
| | scaled (longitude, knot1) | **-0.810026857** | **0.367531771** | **-2.203964179** | **0.027526863** |
| | scaled (longitude, knot2) | **-0.427256011** | **0.110065787** | **-3.881823982** | **0.000103676** |
| | scaled (longitude, knot3) | **-1.154167859** | **0.345034432** | **-3.345080231** | **0.000822588** |
| | scaled (longitude, knot4) | **0.421503109** | **0.10509295** | **4.010764846** | **6.05E-05** |
| | scaled (longitude, knot5) | **-0.307938262** | **0.101744599** | **-3.026580904** | **0.002473366** |
| | SW1 : scaled (residual Length) | **0.211016874** | **0.182101203** | **1.158789018** | **0.246542195** |
| | SW2 : scaled (residual Length) | **0.150328425** | **0.067704366** | **2.220365305** | **0.02639398** |
| | SW3 : scaled (residual Length) | **-0.029296911** | **0.080438579** | **-0.364214676** | **0.715697709** |
| | SW1 : scaled (six6) | **-0.335381813** | **0.17597978** | **-1.905797439** | **0.056676491** |
| | SW2 : scaled (six6) | **-0.166961262** | **0.067581096** | **-2.470532047** | **0.013491223** |
| | SW3 : scaled (six6) | **0.023278062** | **0.099677088** | **0.233534729** | **0.815346205** |
| | SW1 : scaled (vgll3) | **0.181546249** | **0.20157175** | **0.900653237** | **0.36777272** |
| | SW2 : scaled (vgll3) | **-0.037726097** | **0.070513964** | **-0.535015976** | **0.59263881** |
| | SW3 : scaled (vgll3) | **0.049348331** | **0.075071681** | **0.657349478** | **0.510956229** |
| Conditional, zero truncated model | | **Estimate** | **Std. Error** | **z value** | **Pr(>\|z\|)** |
| | SW1 | **0.150661924** | **0.226356201** | **0.665596629** | **0.505668961** |
| | SW2 | **0.288138112** | **0.157949429** | **1.824242821** | **0.068115371** |
| | SW3 | **-0.725434359** | **0.202299167** | **-3.585948335** | **0.000335855** |
| | scaled (sDays1) | **-0.397340364** | **0.197731404** | **-2.009495491** | **0.044484612** |
| | scaled (sDays2) | **-0.530685137** | **0.174436294** | **-3.042286235** | **0.002347885** |
| | scaled (sDays3) | **-0.690898971** | **0.197433099** | **-3.499408033** | **0.000466292** |

| | | | | | |
|---|---|---|---|---|---|
| | scaled (sDays4) | **-0.196327598** | **0.094635617** | **-2.074563509** | **0.038027007** |
| | scaled (sDays5) | **-1.24727363** | **0.294239291** | **-4.238977143** | **2.25E-05** |
| | scaled (sLong1) | **0.24853801** | **0.420439332** | **0.591138819** | **0.554427412** |
| | scaled (sLong2) | **0.146007327** | **0.140390746** | **1.040006776** | **0.298336753** |
| | scaled (sLong3) | **0.707502599** | **0.383182831** | **1.846383869** | **0.064836488** |
| | scaled (sLong4) | **-0.032638553** | **0.11554956** | **-0.282463673** | **0.777587998** |
| | scaled (sLong5) | **-0.237631748** | **0.130376372** | **-1.822659618** | **0.06835496** |
| | SW1 : scaled (residual Length) | **-0.140253923** | **0.185859628** | **-0.754622854** | **0.450475307** |
| | SW2 : scaled (residual Length) | **-0.114095146** | **0.070896233** | **-1.609325918** | **0.107545095** |
| | SW3 : scaled (residual Length) | **-0.534059375** | **0.100447423** | **-5.31680513** | **1.06E-07** |
| | SW1 : scaled (six6) | **0.351847281** | **0.177610348** | **1.981006649** | **0.047590528** |
| | SW2 : scaled (six6) | **0.012834988** | **0.070662442** | **0.181638047** | **0.855866788** |
| | SW3 : scaled (six6) | **-0.031088284** | **0.126405441** | **-0.24594103** | **0.805727882** |
| | SW1 : scaled (vgll3) | **-0.020666622** | **0.187090598** | **-0.110463176** | **0.912042053** |
| | SW2 : scaled (vgll3) | **-0.049415339** | **0.07838416** | **-0.630425064** | **0.528416517** |
| | SW3 : scaled (vgll3) | **-0.008100314** | **0.095934427** | **-0.084435941** | **0.932709833** |
| | Random variance | | | | |
| | population | | | | |
| Zero inflation (binomial) model | **0.00393** | | | | |
| Conditional, zero truncated model | **6.27E-10** | | | | |

**Supp. table 8:** Coefficients of conditional, zero truncated model explaining average prey weight in Atlantic salmon diet. Scaled variables are centered to zero mean, and scaled to one standard deviation.

| | | Estimate | Std. Error | z value | Pr(>\|z\|) |
|---|---|---|---|---|---|
| Zero inflation (binomial) model | SW1 | **-0.445623979** | **0.167708847** | **-2.657128624** | **0.007880937** |
| | SW2 | **-0.074028965** | **0.07276505** | **-1.017369804** | **0.308977541** |
| | SW3 | **0.424101306** | **0.098219136** | **4.317909137** | **1.58E-05** |
| | scaled (days, knot1) | **0.685058379** | **0.157140711** | **4.359521941** | **1.30E-05** |
| | scaled (days, knot2) | **0.775023744** | **0.144473548** | **5.364468116** | **8.12E-08** |
| | scaled (days, knot3) | **0.743665052** | **0.160470319** | **4.634284124** | **3.58E-06** |
| | scaled (days, knot4) | **0.204679154** | **0.079396952** | **2.577922055** | **0.009939642** |
| | scaled (days, knot5) | **1.383924195** | **0.232176728** | **5.960649931** | **2.51E-09** |
| | scaled (longitude, knot1) | **-0.794002253** | **0.368499316** | **-2.15469125** | **0.031186003** |
| | scaled (longitude, knot2) | **-0.422150655** | **0.110282264** | **-3.827910671** | **0.000129236** |
| | scaled (longitude, knot3) | **-1.133569049** | **0.345765743** | **-3.278430764** | **0.00104386** |
| | scaled (longitude, knot4) | **0.415695015** | **0.10480305** | **3.966440066** | **7.30E-05** |
| | scaled (longitude, knot5) | **-0.306369444** | **0.102257574** | **-2.996056256** | **0.00273496** |
| | SW1 : scaled (residual Length) | **0.212031772** | **0.182199601** | **1.163733461** | **0.244532045** |
| | SW2 : scaled (residual Length) | **0.149775011** | **0.067727903** | **2.211422543** | **0.02700659** |
| | SW3 : scaled (residual Length) | **-0.02239038** | **0.080516081** | **-0.278085816** | **0.780946483** |
| | SW1 : scaled (six6) | **-0.334499965** | **0.176036653** | **-1.900172262** | **0.057410517** |
| | SW2 : scaled (six6) | **-0.166103415** | **0.067619921** | **-2.456427219** | **0.014032622** |
| | SW3 : scaled (six6) | **0.028084434** | **0.099784298** | **0.281451433** | **0.778364174** |
| | SW1 : scaled (vgll3) | **0.18216426** | **0.201629238** | **0.903461533** | **0.366280995** |
| | SW2 : scaled (vgll3) | **-0.037402186** | **0.070551676** | **-0.530138868** | **0.596015653** |
| | SW3 : scaled (vgll3) | **0.038442601** | **0.075156195** | **0.511502761** | **0.608999056** |
| Conditional, zero truncated model | | **Estimate** | **Std. Error** | **z value** | **Pr(>\|z\|)** |
| | SW1 | **-0.452942348** | **0.208001101** | **-2.177595914** | **0.029436134** |
| | SW2 | **0.492788724** | **0.117194154** | **4.204891707** | **2.61E-05** |
| | SW3 | **1.643862271** | **0.135112346** | **12.16663255** | **4.68E-34** |
| | scaled (sDays1) | **-0.279961872** | **0.176436858** | **-1.586753898** | **0.112568394** |

| | | | | | |
|---|---|---|---|---|---|
| | scaled (sDays2) | **-0.22626811** | **0.159325791** | **-1.42015997** | **0.155561114** |
| | scaled (sDays3) | **-0.021386999** | **0.175806629** | **-0.121650697** | **0.903175662** |
| | scaled (sDays4) | **-0.093470322** | **0.083652594** | **-1.117363103** | **0.263839104** |
| | scaled (sDays5) | **-0.224628527** | **0.262166775** | **-0.856815387** | **0.391546921** |
| | scaled (sLong1) | **1.517792991** | **0.432936187** | **3.505812261** | **0.000455216** |
| | scaled (sLong2) | **0.552884333** | **0.135925866** | **4.067543197** | **4.75E-05** |
| | scaled (sLong3) | **1.243165866** | **0.393466462** | **3.159521807** | **0.001580283** |
| | scaled (sLong4) | **-0.119629303** | **0.118598632** | **-1.008690406** | **0.313123134** |
| | scaled (sLong5) | **0.349859752** | **0.133097543** | **2.628596627** | **0.008573799** |
| | SW1 : scaled (residual Length) | **-0.305813004** | **0.190226327** | **-1.607627131** | **0.107916867** |
| | SW2 : scaled (residual Length) | **0.2477306** | **0.067705483** | **3.658944447** | **0.000253256** |
| | SW3 : scaled (residual Length) | **0.310762723** | **0.091671019** | **3.389977828** | **0.000698983** |
| | SW1 : scaled (six6) | **0.375974948** | **0.19693412** | **1.909140718** | **0.056243941** |
| | SW2 : scaled (six6) | **0.146190112** | **0.071111304** | **2.055792873** | **0.039802485** |
| | SW3 : scaled (six6) | **-0.094954178** | **0.105954748** | **-0.896176714** | **0.370158391** |
| | SW1 : scaled (vgll3) | **-0.009103335** | **0.182057755** | **-0.050002459** | **0.960120429** |
| | SW2 : scaled (vgll3) | **0.124224948** | **0.074794885** | **1.660874912** | **0.096738572** |
| | SW3 : scaled (vgll3) | **0.083106637** | **0.081605111** | **1.018399901** | **0.308487949** |
| | Random variance | | | | |
| | population | | | | |
| Zero inflation (binomial) model | **0.006575** | | | | |
| Conditional, zero truncated model | **0.12** | | | | |

**Supp. table 9:** Generalized additive prey preference model for each species, measured binomially as the proportional contribution of specific prey species to the total stomach content.

| Species | Linear terms | Estimate | Std. Error | t value | Pr(>\|t\|) |
|---|---|---|---|---|---|
| **San eel** | SW1 | -0.60052478 | 0.2506048 | -2.396302046 | 0.0167551 |
| | SW2 | -0.967555233 | 0.1205907 | -8.023461456 | 3.03E-15 |
| | SW3 | -2.21052119 | 0.2131618 | -10.37015769 | 6.17E-24 |
| | SW1 : scaled (residual Length) | 2.392230717 | 3.5723458 | 0.66965261 | 0.503243 |
| | SW2 : scaled (residual Length) | -3.589414518 | 1.5743045 | -2.280000216 | 0.0228299 |
| | SW3 : scaled (residual Length) | -5.167374501 | 2.465996 | -2.095451262 | 0.0363967 |
| | SW1 : scaled (six6) | -0.045970201 | 0.2442015 | -0.188247027 | 0.8507235 |
| | SW2 : scaled (six6) | 0.021626546 | 0.1081936 | 0.199887471 | 0.8416116 |
| | SW3 : scaled (six6) | -0.063732458 | 0.1937302 | -0.328975272 | 0.7422473 |
| | SW1 : scaled (vgll3) | -0.074990333 | 0.2648959 | -0.283093631 | 0.7771671 |
| | SW2 : scaled (vgll3) | 0.08820433 | 0.111258 | 0.792791241 | 0.4280986 |
| | SW3 : scaled (vgll3) | 0.088429737 | 0.1647431 | 0.536773704 | 0.5915504 |
| | **Smooth terms** | **edf** | **Ref.df** | **F** | **p-value** |
| | scaled(DAY) | 2.993901503 | 2.9939015 | 6.095705208 | 0.0004573 |
| | scaled(long) | 6.758485905 | 6.7584859 | 9.595347964 | 3.50E-11 |
| | population | 2.08E-05 | 123 | 1.73E-07 | 0.4443376 |
| **Capelin** | SW1 | -0.789639186 | 0.2519124 | -3.134578837 | 0.0017738 |
| | SW2 | -1.346177895 | 0.1263673 | -10.65289929 | 4.13E-25 |
| | SW3 | -2.104510635 | 0.2195978 | -9.583479308 | 7.91E-21 |
| | SW1 : scaled (residual Length) | -2.607704634 | 3.5657515 | -0.731319789 | 0.4647641 |
| | SW2 : scaled (residual Length) | 1.020833231 | 1.6101771 | 0.633988154 | 0.5262411 |
| | SW3 : scaled (residual Length) | -3.627072741 | 2.4637857 | -1.472154318 | 0.1413102 |
| | SW1 : scaled (six6) | -0.060613787 | 0.2530223 | -0.239559059 | 0.8107238 |
| | SW2 : scaled (six6) | -0.180947939 | 0.114276 | -1.583429028 | 0.1136564 |
| | SW3 : scaled (six6) | 0.341408886 | 0.2242019 | 1.522774283 | 0.1281478 |
| | SW1 : scaled (vgll3) | 0.083132346 | 0.2682153 | 0.309946328 | 0.7566697 |

| | | | | | |
|---|---|---|---|---|---|
| | SW2 : scaled (vgll3) | -0.20066241 | 0.1179432 | -1.701348346 | 0.0892046 |
| | SW3 : scaled (vgll3) | -0.297694647 | 0.1675406 | -1.776850297 | 0.0759126 |
| | **Smooth terms** | **edf** | **Ref.df** | **F** | **p-value** |
| | scaled(DAY) | 2.502146859 | 2.5021469 | 7.242203336 | 0.0001875 |
| | scaled(long) | 1.000010194 | 1.0000102 | 51.77249193 | 1.22E-12 |
| | population | 7.10E-09 | 124 | 4.10E-11 | 0.9310087 |
| **Herring** | SW1 | -1.391853747 | 0.2653217 | -5.245910374 | 1.92E-07 |
| | SW2 | -0.486569989 | 0.1188431 | -4.094223175 | 4.60E-05 |
| | SW3 | 0.698391314 | 0.1684402 | 4.146228111 | 3.68E-05 |
| | SW1 : scaled (residual Length) | 6.643836033 | 4.1380745 | 1.605538046 | 0.108708 |
| | SW2 : scaled (residual Length) | 2.141664672 | 1.6068553 | 1.33282981 | 0.1829074 |
| | SW3 : scaled (residual Length) | 3.760154212 | 2.1238249 | 1.770463384 | 0.0769708 |
| | SW1 : scaled (six6) | -0.075850003 | 0.2839296 | -0.267143735 | 0.7894165 |
| | SW2 : scaled (six6) | 0.178710274 | 0.1154384 | 1.548101348 | 0.1219311 |
| | SW3 : scaled (six6) | -0.241836855 | 0.1729131 | -1.39860361 | 0.1622584 |
| | SW1 : scaled (vgll3) | -0.098199684 | 0.2850859 | -0.344456428 | 0.7305793 |
| | SW2 : scaled (vgll3) | 0.003670978 | 0.1212537 | 0.030275176 | 0.975854 |
| | SW3 : scaled (vgll3) | 0.07133899 | 0.1346883 | 0.529659799 | 0.5964716 |
| | **Smooth terms** | **edf** | **Ref.df** | **F** | **p-value** |
| | scaled(DAY) | 1.000002543 | 1.0000025 | 72.70853915 | 5.31E-17 |
| | scaled(long) | 3.537911425 | 3.5379114 | 42.03594295 | 1.40E-28 |
| | population | 5.77E-06 | 124 | 4.66E-08 | 0.4800687 |
| **Haddock** | SW1 | -41.18627138 | 5187.0694 | -0.007940181 | 0.9936664 |
| | SW2 | -3.619308356 | 0.2938553 | -12.31663539 | 1.95E-32 |
| | SW3 | -2.387135937 | 0.2662801 | -8.964756853 | 1.63E-18 |
| | SW1 : scaled (residual Length) | -21.07580971 | 24.086287 | -0.875012822 | 0.3817892 |
| | SW2 : scaled (residual Length) | 7.228440165 | 3.4410701 | 2.100637316 | 0.0359372 |
| | SW3 : scaled (residual Length) | 3.846619807 | 2.7760192 | 1.385660384 | 0.1661775 |
| | SW1 : scaled (six6) | -0.577679805 | 3.006245 | -0.192159923 | 0.8476582 |

| | | | | | |
|---|---|---|---|---|---|
| | SW2 : scaled (six6) | 0.115827182 | 0.2509622 | 0.461532301 | 0.6445229 |
| | SW3 : scaled (six6) | 0.106126108 | 0.228151 | 0.465157268 | 0.641926 |
| | SW1 : scaled (vgll3) | 24.20962992 | 3279.8003 | 0.007381434 | 0.9941121 |
| | SW2 : scaled (vgll3) | 0.265665408 | 0.2326657 | 1.141833079 | 0.2538126 |
| | SW3 : scaled (vgll3) | 0.057011782 | 0.1740237 | 0.327609347 | 0.7432796 |
| | **Smooth terms** | **edf** | **Ref.df** | **F** | **p-value** |
| | scaled(DAY) | 1.000000193 | 1.0000002 | 11.33308323 | 0.0007914 |
| | scaled(long) | 2.479590964 | 2.479591 | 8.868644537 | 3.89E-05 |
| | population | 6.29744996 | 122 | 0.068027213 | 0.1236614 |

**Supp. table 10:** Generalized additive prey preference model for each species, measured binomially as the proportional contribution of specific prey species to the total stomach content. The model does not include sea age and residual length as terms.

| Species | Linear terms | Estimate | Std. Error | t value | Pr(>\|t\|) | | Smooth terms | edf | Ref.df | F | p-value |
|---|---|---|---|---|---|---|---|---|---|---|---|
| Sand eel | (Intercept) | -1.235184007 | 0.0968776 | -12.74993853 | 1.61E-34 | | s(scale(DAY)) | 1.0000001 | 1.0000001 | 2.4591581 | 0.1171644 |
| | scale(six6) | -0.203780725 | 0.0774587 | -2.630831104 | 0.008653509 | | s(scale(long)) | 6.7780124 | 6.7780124 | 10.365614 | 2.75E-12 |
| | scale(vgll3.imp) | -0.069140332 | 0.079336 | -0.871486981 | 0.383705252 | | s(popF) | 5.0285513 | 123 | 0.0479374 | 0.2045599 |
| Capelin | (Intercept) | -1.443955425 | 0.0915057 | -15.77994431 | 4.04E-50 | | s(scale(DAY)) | 2.8893904 | 2.8893904 | 5.271529 | 0.0009898 |
| | scale(six6) | -0.135764548 | 0.0808817 | -1.678557153 | 0.093559444 | | s(scale(long)) | 1.0000003 | 1.0000003 | 59.280693 | 3.19E-14 |
| | scale(vgll3.imp) | -0.25869961 | 0.0850299 | -3.042454412 | 0.00240953 | | s(popF) | 4.26E-07 | 123 | 2.05E-09 | 0.9895952 |
| | (Intercept) | -0.236774242 | 0.0807378 | -2.932631148 | 0.00344005 | | s(scale(DAY)) | 1.0000001 | 1.0000001 | 48.032595 | 7.43E-12 |
| Herring | scale(six6) | 0.234629433 | 0.079414 | 2.95450946 | 0.003207329 | | s(scale(long)) | 3.1520614 | 3.1520614 | 48.556121 | 2.29E-30 |
| | scale(vgll3.imp) | 0.168787526 | 0.0777616 | 2.170577678 | 0.030205106 | | s(popF) | 2.8933842 | 123 | 0.0269057 | 0.2479188 |
| | (Intercept) | -3.110628396 | 0.1941955 | -16.01802625 | 2.17E-51 | | s(scale(DAY)) | 1.0000001 | 1.0000001 | 21.483272 | 4.04E-06 |
| | scale(six6) | 0.345592954 | 0.1485126 | 2.327027182 | 0.020169739 | | s(scale(long)) | 2.4853189 | 2.4853189 | 9.3271825 | 2.75E-05 |
| Haddock | scale(vgll3.imp) | 0.304559899 | 0.1306665 | 2.330818753 | 0.019967928 | | s(popF) | 7.4189751 | 123 | 0.0805933 | 0.0868028 |

**Supp. table 11:** Coefficients of the model explaining sampling date in relation to sea age, and life history genomic regions (*six6* and *vgll3*), as well as smoother functions of longitude and population. Scaled variables are centered to zero mean, and scaled to one standard deviation. Modelling was performed using *gam* function in *mgcv* package. Response variable was modelled as binomial, where success was equal to number of days to avoiding sampling in the range of date of sampling.

| **Linear terms** | **Estimate** | **Std. Error** | **t value** | **Pr(>\|t\|)** |
|---|---|---|---|---|
| SW1 | 3.81805474 | 0.083224754 | 45.8764315 | 0.00E+00 |
| SW2 | 3.460917804 | 0.045199595 | 76.56966439 | 0 |
| SW3 | 3.090860656 | 0.054935672 | 56.263272 | 0 |
| SW1 : scaled (six6) | -0.017535338 | 0.080560674 | -0.217666226 | 0.827710955 |
| SW2 : scaled (six6) | 0.099070491 | 0.031739773 | 3.121335828 | 0.001825557 |
| SW3 : scaled (six6) | 0.094992467 | 0.048410135 | 1.962243385 | 0.049869954 |
| SW1 : scaled (vgll3) | 0.045985521 | 0.094132917 | 0.488516906 | 0.625236278 |
| SW2 : scaled (vgll3) | -0.035381104 | 0.035383065 | -0.999944587 | 0.317455954 |
| SW3 : scaled (vgll3) | 0.014540231 | 0.035979371 | 0.40412687 | 0.686161802 |
| **Smooth terms** | **edf** | **Ref.df** | **F** | **p-value** |
| scale(long) | 1.000000082 | 1.000000082 | 4.332845323 | 0.037505811 |
| population | 41.63729405 | 133 | 0.887874138 | 2.16E-14 |
